# Unwinding of RNA G-quadruplexes induces mouse and human totipotency

**DOI:** 10.64898/2026.02.13.703305

**Authors:** Yicong Dai, Xucong Teng, Qiushuang Zhang, Ronghui Li, Yuncong Wu, Difei Hu, Xiangdong Zhang, Yuhan Liu, Wenxuan Hu, Yeyin Li, Xin Meng, Na Zhang, Xuena Cao, Lili Dong, Chengjia Mi, Jing Zhang, Jiang Xie, Kun Qu, Juan Carlos Izpisua Belmonte, Haisong Liu, Jinghong Li

**Author notes:** Yicong Dai, Xucong Teng and Qiushuang Zhang contributed equally to this work.

## Abstract

Totipotent stem cells (TotiSCs) have significant application prospects in regenerative medicine and assisted reproductive technologies. The mRNAs of totipotency-related genes are rich in G-quadruplex (G4) structures, while the G4 density of pluripotency-related genes mRNAs is relatively low, suggesting a potential functional link between RNA G4s and the totipotent state. Nevertheless, how G4s are involved in the establishment or maintenance of mouse and human totipotency remains poorly understood. Here, we demonstrate that FC3, a small-molecule RNA G4 unwinder, enables efficient induction and maintenance of mouse and human TotiSCs (mG4TotiSCs and hG4TotiSCs). These FC3-induced TotiSCs closely recapitulate the molecular features of 2-cell-stage blastomeres via transcriptomic and epigenomic profiling. Functionally, both mG4TotiSCs and hG4TotiSCs exhibit totipotent capacity: they differentiate into embryonic and extraembryonic lineages in vitro and in chimeric embryos; moreover, they autonomously self-organize into blastocyst-like structures without exogenous signalling. Mechanistically, we identify that Gata2 mRNA, a G4-containing transcript, is essential for FC3-mediated reprogramming. Genetic disruption of Gata2 would abolish totipotency induction. Collectively, RNA G4 unwinding is a new regulatory axis governing totipotency acquisition and maintenance, providing insight into the functional interplay between nucleic acid secondary structures and cell fate plasticity.

## Introduction

Totipotent stem cells (TotiSCs) possess the unique capacity to differentiate into both embryonic and extraembryonic lineages. In vivo, this capability is naturally restricted to the zygote and blastomeres of the 2-cell to 4-cell stage, which are universally recognized as the sole endogenous sources of totipotency^1^. The derivation and maintenance of TotiSCs from non-germline somatic or pluripotent cells is crucial in stem cell biology, with broad implications for regenerative medicine, assisted reproductive technologies, and mechanistic studies of early human development^2^. To date, multiple strategies have been reported to reprogram mouse embryonic stem cells (mESCs), which are pluripotent cells derived from the inner cell mass (ICM), into a totipotent state^3–14^. Early approaches mainly relied on genetic interventions, such as targeted knockout or overexpression of totipotency-associated factors; however, these methods entail technical complexity, low efficiency, and inherent risks of genomic instability. Comparatively, chemical reprogramming, achieved by supplementing defined small molecules into culture media without genetic modification, offers a more precise, scalable, and clinically translatable strategy to modulate cellular plasticity. While recent studies have employed multi-component small-molecule cocktails to induce totipotency in mESCs, the concurrent modulation of diverse signalling and epigenetic pathways has hindered the clarification of core drivers of the pluripotency-to-totipotency transition. Whatmore, for human cells, only a handful of chemically reprogramed totipotent-like cells have been reported^15–17^, including 8-cell-like cells (8CLCs)^17^ and human totipotent blastomere-like cells (hTBLCs)^15^. Consequently, the feasibility of achieving robust totipotency induction in mouse and human cells, and the extent to which conserved regulatory principles govern this transition across species, still remains an open and pivotal question.

G-quadruplexes (G4s) are special nucleic acid secondary structures formed through Hoogsteen hydrogen bonding among four guanine residues in G-rich sequences^18^, which can regulate early embryo development and stem cell differentiation^19^ **(Fig. 1a).** They are evolutionarily conserved and widely distributed across mammalian genomes and transcriptomes, occurring in both genomic DNA (DNA G4s or dG4s) and RNA (RNA G4s or rG4s). Functionally, G4s serve as dynamic regulatory elements that modulate essential cellular processes, including DNA replication, transcriptional initiation and elongation, RNA splicing, nuclear export, and translation, which are exquisitely coordinated during early embryogenesis and stem cell fate transitions. Accumulating evidence indicates that G4 dysregulation contributes to the pathogenesis of multiple human diseases, notably cancer^20^, neurodegenerative disorders^21^, and viral infections^22^, positioning G4s as promising therapeutic targets^23–25^. Correspondingly, a growing number of selective small-molecule ligands, acting either as G4 stabilizers or unwinders, have been developed to probe or manipulate G4 structure and function in biological contexts^26^. Despite these advances, the concrete mechanism of how G4-targeted chemical modulation control stem cell reprogramming dynamics and pluripotency-to-totipotency transitions remains largely uninvestigated.

**Fig. 1.**
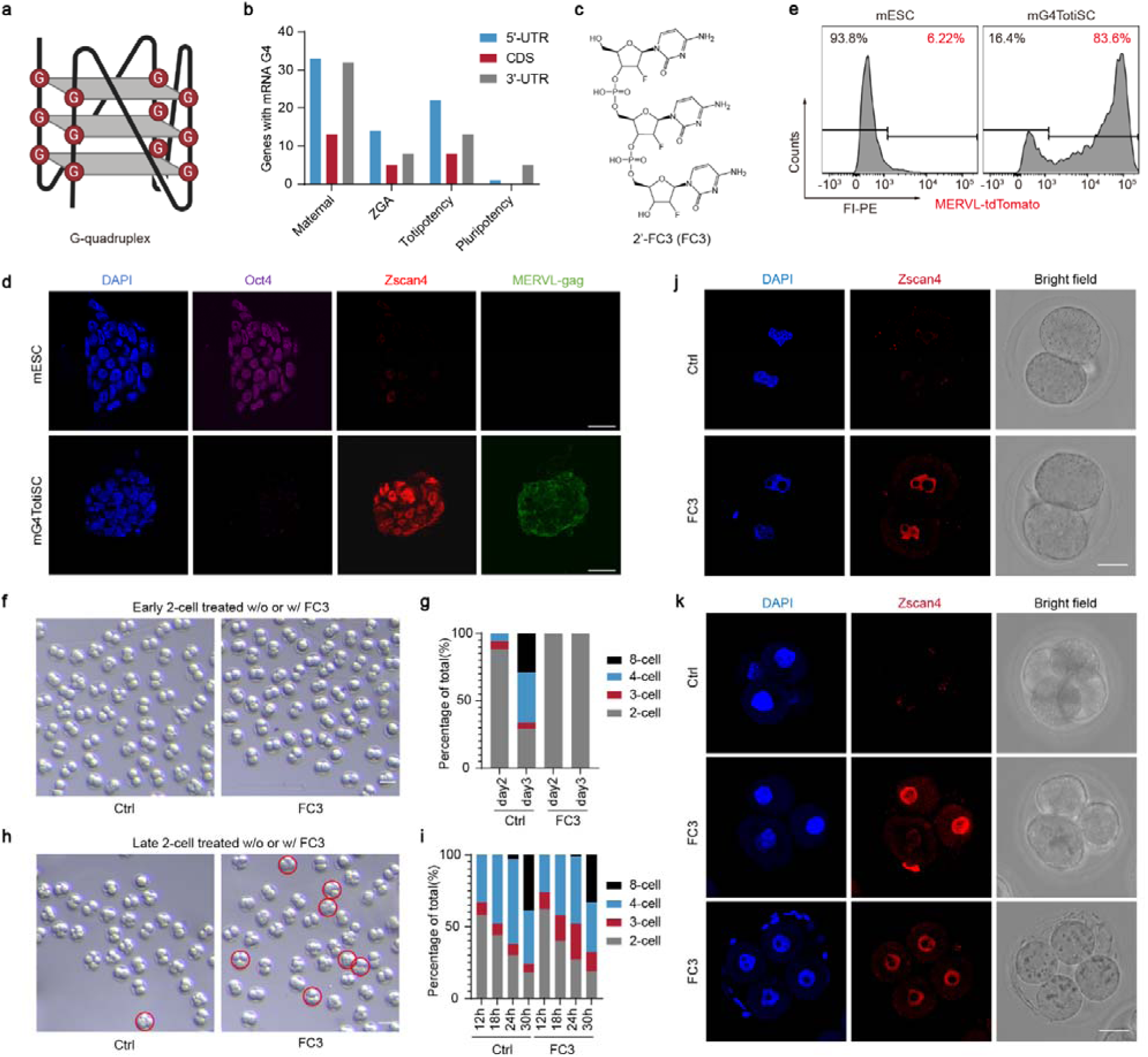
rG4 unwinding by FC3 enables the establishment and maintenance of totipotent gene expression. **a**, Schematic diagram of a G4 structure. **b**, Numbers of mouse maternal genes, ZGA genes, totipotency genes and pluripotency genes with a G4 structure in the corresponding mRNAs. **c**, Chemical structure of 2’-F C3 (FC3). **d**, Representative immunofluorescence images of Oct4, Zscan4 and MERVL-gag in mESCs or mG4TotiSCs. Scale bar, 25 μm. **e**, FACS analysis of MERVL-tdTomato^+^ mESCs or mG4TotiSCs. **f**, Imaging of the arrest of mouse early 2-cell embryos by FC3. Early 2-cell embryos were treated without (Ctrl group) or with (FC3 group) FC3. The embryo morphology at day 2 after treatment of FC3 was shown in the bright field pictures. Scale bar, 100 μm. **g**, Percentage of embryos reaching different developmental stages at day 2, day 3 post treatment of mouse early 2-cell embryos with FC3. **h**, Imaging of the arrest of mouse late 2-cell embryos by FC3. Late 2-cell embryos were treated without (Ctrl group) or with (FC3 group) FC3. The embryo morphology at 24 h after treatment of FC3 was shown in the bright field pictures. Scale bar, 100 μm. **i**, Percentage of embryos reaching different development stages at 12 h, 18 h, 24 h, 30 h after treatment of mouse late 2-cell embryos with FC3. **j**, Immunofluorescence images of Zscan4 in mouse early 2-cell embryos treated without (Ctrl group) or with (FC3 group) FC3 after 24 h. Scale bar, 30 μm. **k**, Immunofluorescence images of Zscan4 in FC3-treated mouse embryos. Late 2-cell embryos were treated without (Ctrl group) or with (FC3 group) FC3 for 24 h. 4-cell embryos or 3-cell embryos were observed at this time point and immunostaining of Zscan4 was performed. Scale bar, 30 μm.

Here, we systematically investigated the functional involvement of rG4s in totipotency acquisition. We demonstrate that chemical unwinding of rG4 structures represents a previously unidentified regulatory mechanism capable of driving the pluripotency-to-totipotency transition in mouse and human embryonic stem cells (mESCs and hESCs). Treatment of mESCs or hESCs with FC3, a small-molecule rG4 unwinder, enabled the robust derivation of totipotent stem cells (termed mG4TotiSCs and hG4TotiSCs) that fulfill stringent molecular, epigenetic, and functional criteria for totipotency. Specifically, mG4TotiSCs and hG4TotiSCs exhibit stable self-renewal under reproducible culture conditions supplemented with FC3, as well as transcriptional and epigenomic profiles and bona fide developmental competence highly similar to those of native 2-cell-stage blastomeres: they contribute extensively to both embryonic and extraembryonic tissues in chimeric mouse embryos, as well as autonomously self-organize into blastocyst-like structures without exogenous signalling. Mechanistically, FC3-mediated reprogramming depends critically on the unwinding of a conserved rG4 motif within the 5′-UTR of mouse and human GATA2 mRNAs, leading to enhanced GATA2 protein synthesis and subsequent activation of a downstream totipotency-associated gene network. To summarize, our work establishes rG4 structures as a direct, druggable regulator of cell fate plasticity during early mammalian development, and provides the key experimental evidence linking rG4 structural remodelling to totipotency regulation in stem cell biology.

## Results

### G4 unwinding induces mouse totipotency

Maternal genes, zygotic genome activation (ZGA) genes, totipotency genes, and pluripotency genes collectively regulate critical events in gametogenesis and early embryogenesis^3,4^. To explore a potential regulatory role for rG4s in this context, we performed a systematic in silico analysis of rG4-forming sequences within the mRNA transcripts of these functionally defined gene sets. Using pqsfinder^27^, an exhaustive and imperfection-tolerant search tool for potential quadruplex-forming sequences in R, we scanned the full-length transcriptome of the mouse reference genome (mm39) to identify all high-confidence rG4 motifs. A minimum G-score threshold of 82 was applied to prioritize motifs with strong thermodynamic propensity for stable rG4 formation. We further annotated the genomic localization of each predicted rG4 motif across canonical mRNA regions: the 5′-untranslated region (5′-UTR), coding sequence (CDS), and 3′-untranslated region (3′-UTR). Strikingly, among the 59 pluripotency genes, only six harbored predicted rG4 motifs, exclusively in the 5′-UTR (n = 1) or 3′-UTR (n = 5), whereas maternal, ZGA, and totipotency genes exhibited significantly higher rG4 motif density, particularly within CDS regions (**Fig. 1b**). These transcriptome-wide disparities prompted the hypothesis that rG4 structural dynamics may serve as a previously underappreciated molecular switch governing the transition between totipotent and pluripotent states during early mammalian development, and that targeted modulation of rG4s could represent a viable strategy for reprogramming mESCs toward expanded developmental potency.

To experimentally test this hypothesis, we assessed whether chemical modulation of endogenous rG4s in mouse embryonic stem cells (mESCs) could drive reprogramming toward a totipotent state. We first conducted a systematic comparative screen of established G4-targeting compounds, including both stabilizers and unwinders, evaluating their capacity to induce molecular signatures of totipotency. We selected four well-characterized G4 stabilizers from previous research: pyridostatin (PDS)^28^, PhenDC3^29^, RGB-1^30^, and TMPyP4^31^ (**Extended Data Fig. 1a**). In parallel, given the scarcity of rigorously validated small-molecule rG4 unwinders^32^, we prioritized two candidates with robust biochemical and cellular activity: PhpC^33^ and 2′-F-C3 (FC3)^34^, both of which demonstrate high-efficiency rG4 resolution in vitro and functional target engagement in vivo (**Fig. 1c; Extended Data Fig. 1a**). To enable sensitive and real-time detection of totipotency acquisition, we generated two mESC reporter lines (**Extended Data Fig. 1b**): (i) MERVL-TdTomato, in which TdTomato expression is driven by MERVL long terminal repeat (LTR) elements, endogenous enhancers specifically active in the 2-cell stage and canonical totipotent states^35,36^; and (ii) OCT4-EGFP, in which EGFP is under the control of the OCT4 promoter, a locus whose transcriptional activation is a hallmark of pluripotency^4,37^. Dual-reporter readouts thus provide orthogonal, functionally anchored metrics for totipotency induction. Immunofluorescence analysis confirmed the expected expression profile in both reporter lines: robust nuclear OCT4 protein expression in pluripotent mESCs and absence of ZSCAN4, a canonical totipotency-associated protein, in the undifferentiated state (**Extended Data Fig. 1c, d**). Before totipotency induction assays, we systematically evaluated compound cytotoxicity by exposing mESCs cultured on mitotically inactivated mouse embryonic fibroblasts (MEFs) in serum/LIF medium to a concentration gradient (0.5–10 μM) of each G4 modulator. Two rG4 unwinders, FC3 and PhpC, exhibited minimal impact on mESC proliferation across the tested range, whereas all four G4 stabilizers induced dose-dependent suppression of cell growth, with significant inhibition observed above 3 μM (**Extended Data Fig. 1e, f**). We next assessed functional reprogramming capacity by quantifying MERVL-driven TdTomato activation in MERVL-TdTomato mESCs. Strikingly, treatment with 5 μM FC3 yielded the highest proportion of TdTomato-positive cells (∼12%), closely paralleled by PhpC; in contrast, none of the G4 stabilizers elicited detectable MERVL activation (**Extended Data Fig. 1g**). Collectively, these data demonstrate that disruption of rG4 structures is sufficient to initiate molecular reprogramming toward a totipotent-like state.

Next, we use a combinatorial screening approach to systematically optimized the culture conditions for efficient totipotency induction, including basal medium formulation, auxiliary small-molecule supplementation, and extracellular matrix support (**Extended Data Fig. 1h–j**). From the optimization, we identified a simple, FC3-based chemical protocol in standard culture conditions, designated SLFM (serum/LIF medium supplemented with 5 μM FC3 and cultured on Matrigel) as optimal for robust and reproducible reprogramming. Under SLFM, mESCs exhibited pronounced transcriptional activation of canonical totipotency-associated genes, including Zscan4d, Zscan4f, Btg2, Mdm2, Zfp352, and Sp110, as quantified by RT–qPCR (**Extended Data Fig. 1k**). Immunofluorescence analysis further confirmed coordinated protein-level remodelling: nuclear OCT4 expression was markedly downregulated, while ZSCAN4 and MERVL-gag proteins, normally undetectable in pluripotent mESCs, was obviously induced (**Fig. 1d**). Flow cytometry revealed that >80% of SLFM-treated cells activated the MERVL-driven TdTomato reporter (**Fig. 1e**), reflecting a high efficiency of functional totipotency induction. Collectively, SLFM is established as an effective platform for generating TotiSCs from pluripotent precursors. The resulting cells were therefore designated mouse G4-unwinder–induced totipotent stem cells (mG4TotiSCs).

We next investigated the impact of G4 unwinding by FC3 on preimplantation mouse embryos, specifically focusing on 2-cell-stage blastomeres. Early and late 2-cell embryos were isolated from ICR mice and cultured in KSOM medium supplemented with 5 μM FC3. Strikingly, FC3 treatment induced complete developmental arrest in 100% of early two-cell embryos (**Fig. 1f-g**), preventing progression beyond the 2-cell stage. Late 2-cell embryos exhibited similar but different developmental dynamics under FC3 treatment (**Fig. 1h-i**): approximately 20% of the cells undergo asymmetric division to form a three-cell embryo, which occurs with a very low probability under standard KSOM culture conditions^38^. Notably, ZSCAN4, whose transcriptional activity is tightly restricted to the two-cell stage and serves as a molecular signature of totipotency, normally becomes undetectable by the late two-cell stage. Immunofluorescence analysis revealed that FC3 significantly extended the expression of ZSCAN4 protein (**Fig. 1j-k**), sustaining nuclear ZSCAN4 signal not only in arrested late two-cell embryos but also in FC3-induced three-cell and even four-cell embryos. These findings demonstrate that chemical unwinding of rG4s preserves and prolongs the expression of core totipotency determinants during early embryogenesis, thereby reinforcing the totipotent state beyond its endogenous temporal window.

### mG4TotiSCs resemble mouse 2-cell blastomeres at both the transcriptomic and epigenetic levels

To assess the long-term expandability and genomic stability of mG4TotiSCs, we evaluated their capacity to sustain self-renewal and molecular identity across serial passaging. After 10 consecutive passages under SLFM conditions, mG4TotiSCs maintained compact, dome-shaped colonies without morphological signs of differentiation or elevated apoptosis (**Extended Data Fig. 2a, b**). Karyotypic analysis confirmed euploid diploid karyotypes at passage 0, 6, and 10 (P0, P6 and P10), with no evidence of aneuploidy, translocations, or other gross chromosomal aberrations (**Extended Data Fig. 2c**). Critically, the defining functional signatures of totipotency, MERVL-LTR-driven reporter activation and concomitant silencing of the OCT4 promoter, remained stably through at least P10 (**Extended Data Fig. 2d**). These data demonstrate that SLFM supports prolonged proliferation with faithful preservation of the totipotent molecular state across multiple generations, proving that mG4TotiSCs is a genetically stable, scalable in vitro model of totipotency.

To comprehensively characterize transcriptional reprogramming, we performed bulk RNA sequencing (RNA-seq) across biological replicates. Relative to control mESCs, mG4TotiSCs exhibited upregulation of canonical totipotency-associated transcripts, including members of the Zscan4 family, Zfp352, Mdm2, and MERVL-derived elements (MERVL-int and MT2_mm), with significant downregulation of core pluripotency factors (Zfp42, Pou5f1/Oct4, Nanog, and Sox2) (**Fig. 2a-b**). Critically, this transcriptional signature was stably maintained through at least P6 and P10, confirming its resilience to prolonged in vitro expansion (**Extended Data Fig. 2e–h**). Parallel quantitative proteomic profiling corroborated these transcriptomic changes, revealing concordant upregulation of totipotency-related proteins and downregulation of OCT4, SOX2 and NANOG (**Fig. 2c**); although inherent limitations in proteomic depth resulted in fewer quantified totipotency and pluripotency-associated proteins, the directional consistency with RNA-seq strongly supports molecular coherence.

**Fig. 2.**
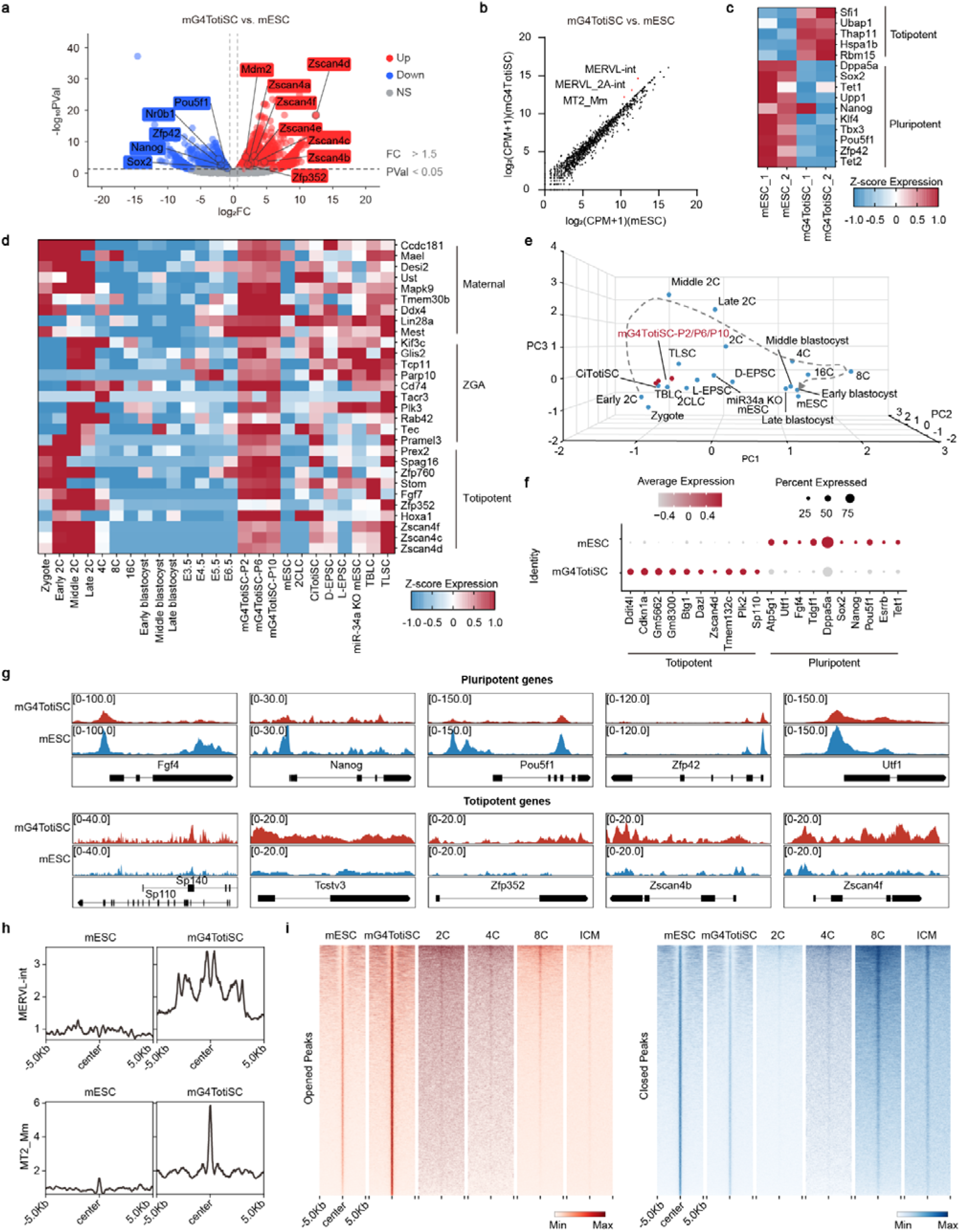
Transcriptomic and epigenomic evaluation of the totipotent characteristics of mG4TotiSCs. **a**, RNA-seq analysis of mESCs and mG4TotiSCs. The global transcriptomic changes are displayed in the volcano plot. Red and blue dots indicate upregulated (fold change D 1.5) and downregulated (fold change D 0.67) genes, with a p-value D 0.05. **b**, Scatterplots displaying the transposon transcript comparison of mESCs and mG4TotiSCs via RNA-seq. Red dots represent the key transposons MERVL-int, MERVL_2A-int, and MT2_Mm. **c**, Heatmap showing the relative protein levels of representative pluripotent and totipotent genes in mESCs and mG4TotiSCs. **d,** Heatmap showing the normalized expression of maternal, ZGA and totipotent genes by RNA-seq of mESCs; mG4TotiSCs (2, 6 or 10 passages); 2CLCs; CiTotiSCs; D-EPSCs; L-EPSCs; miR-24a KO mESCs; TBLCs; TLSCs; and mouse embryos. **f**, Transcriptome-based PCA of mESCs, mG4TotiSCs, 2CLCs, CiTotiSCs, D-EPSCs, L-EPSCs, miR-24a KO mESCs, TBLCs, TLSCs and mouse embryos. **g**, Bubble plot from scRNA-seq showing the relative expression of representative pluripotent and totipotent genes in the mESC and mG4TotiSC clusters. **i**, ATAC-seq peaks around representative pluripotent and totipotent genes in mG4TotiSCs and mESCs. **h**, Line plots showing the relative ATAC-seq signal enrichment in the 5-kb region around the whole-genome-wide MERVL-int and MT2_Mm transposon loci in mESCs and mG4TotiSCs. **i**, Heatmaps showing the opened or closed peaks around mESCs, mG4TotiSCs, 2-cell, 4-cell, and 8-cell blastomeres and the ICM via ATAC-seq; ATAC-seq data for 2-, 4-, and 8-cell blastomeres and the ICM were obtained from a published resource (GEO Accession, GSE66581).

To contextualize mG4TotiSCs within the broader landscape of totipotency models, we conducted an integrative transcriptomic comparison encompassing: (i) mESCs; (ii) in vivo mouse embryos spanning zygote to E6.5; and (iii) previously reported totipotent-like stem cell lines, including 2-cell-like cells (2CLCs) ^36^, chemically induced totipotent stem cells (CiTotiSCs) ^3^, D-EPSCs^6^, L-EPSCs^7^, miR-34a-knockout ESCs^39^, TBLCs^4^, and totipotent-like stem cells (TLSCs) ^8^. In addition to the classic Zscan4 family genes, the maternal genes and ZGA genes specific to zygotes and 2-cell blastomeres, which are also common feature genes of other reported totipotent stem cells, were obviously upregulated in mG4TotiSCs **(Fig. 2d and Extended Data Fig. 2i)**. Furthermore, gene set enrichment analysis (GSEA) of stage-specific transcriptomic signatures from early mouse embryos showed thatmG4TotiSCs exhibited significant positive enrichment in the zygote, 2-cell, and 4-cell gene sets (**Extended Data Fig. 2j**), while the 8-cell and 16-cell gene sets displayed negative enrichment scores. On this basis, unsupervised hierarchical clustering and principal component analysis (PCA) of bulk transcriptomes indicated that mG4TotiSCs consistently clustered with 2-cell blastomeres and showed high transcriptional similarity to established induced totipotent stem cell lines, including TLSCs and CiTotiSCs (**Fig. 2e and Extended Data Fig. 2k**), supporting their strong ability to transition from pluripotency to a totipotent-like state. To resolve cellular heterogeneity at single-cell resolution, we performed single-cell RNA sequencing (scRNA-seq) on mG4TotiSCs and control mESCs. Uniform manifold approximation and projection (UMAP)-based graph clustering clearly segregated the two populations into distinct clusters corresponding to mESCs and mG4TotiSCs. Moreover, supervised scoring using a defined panel of 10 canonical pluripotency-associated genes and 10 totipotency-associated genes enabled unambiguous discrimination between the two cell types (**Fig. 2f**). Notably, mG4TotiSCs exhibited significant downregulation of pluripotency genes and concomitant upregulation of totipotency genes compared to mESCs. Collectively, these integrated transcriptomic analyses provide convergent evidence that mG4TotiSCs closely recapitulate the molecular identity of mouse 2-cell blastomeres.

The transition from pluripotent stem cells to TotiSCs is accompanied by extensive reorganization of the chromatin epigenetic landscape, particularly in chromatin accessibility landscape. To comprehensively map chromatin accessibility changes in mG4TotiSCs, we performed an assay involving transposase-accessible chromatin sequencing (ATAC-seq). As expected, the regulatory regions of core totipotency-associated genes (Sp110, Tcstv3, Zfp352, Zscan4b, and Zscan4f) exhibited significantly increased accessibility in mG4TotiSCs, whereas loci associated with key pluripotency factors (Fgf4, Nanog, Pou5f1, Zfp42, and Utf1) showed pronounced reductions in accessibility (**Fig. 2g**). Importantly, MERVL-derived repetitive elements, specifically MERVL-int and MT2-Mm, also displayed markedly enhanced chromatin accessibility in mG4TotiSCs, consistent with their transcriptional upregulation relative to mESCs (**Fig. 2h**). Genome-wide differential peak analysis revealed that, compared with mESCs, mG4TotiSCs gained 11,553 and lost 18,571 ATAC-seq peaks; these differentially accessible regions showed strong overlap with the accessibility signatures of 2-cell and 4-cell blastomeres (**Fig. 2i**). Together, these data demonstrate that mG4TotiSCs undergo large-scale, developmentally coordinated chromatin remodelling during the pluripotency-to-totipotency transition, thereby acquiring a chromatin accessibility architecture highly analogous to that of early embryonic blastomeres.

Collectively, the results presented in this subsection suggest that mG4TotiSCs generated in vitro are closely similar to totipotent 2-cell blastomeres in vivo at both the transcriptomic and epigenomic levels.

### mG4TotiSCs can differentiate into pluripotent stem cells and extraembryonic lineages in vitro

During early embryonic development, totipotent zygotes, along with 2- and 4-cell blastomeres, gradually differentiate into pluripotent cells. This similar differentiation potential is a crucial characteristic of induced TotiSCs. Therefore, we aimed to determine whether mG4TotiSCs can revert to the pluripotent state. For this purpose, we employed the abovementioned OCT4-EGFP reporter mESCs, which express EGFP in their pluripotent state, to monitor the reversible transition from pluripotency to totipotency. As expected, we found that most OCT4-EGFP-derived mG4TotiSCs lost EGFP expression during FC3 treatment; however, when we replaced the SLFM medium with one lacking FC3, EGFP expression was reactivated in more than 90% of the cells (referred to herein as reversed ESCs [rESCs]) **(Fig. 3a-c)**. Further heatmap analysis of the RNA-seq data revealed that the expression of pluripotency genes was restored in P1 and P2 rESCs **(Fig. 3d)**, whereas totipotency genes and MERVL transcripts were no longer expressed **(Extended Data Fig. 3a, b)**. At the whole-transcriptome level, in P1 and P2 rESCs, the expression of 2782 upregulated genes and 1329 downregulated mG4TotiSC-specific genes returned to levels similar to those in mESCs **(Extended Data Fig. 3c)**. These results indicate that mG4TotiSCs are capable of redifferentiating toward pluripotency and that the transition between pluripotent and totipotent states can be influenced by a single small molecule, namely, FC3.

**Fig. 3.**
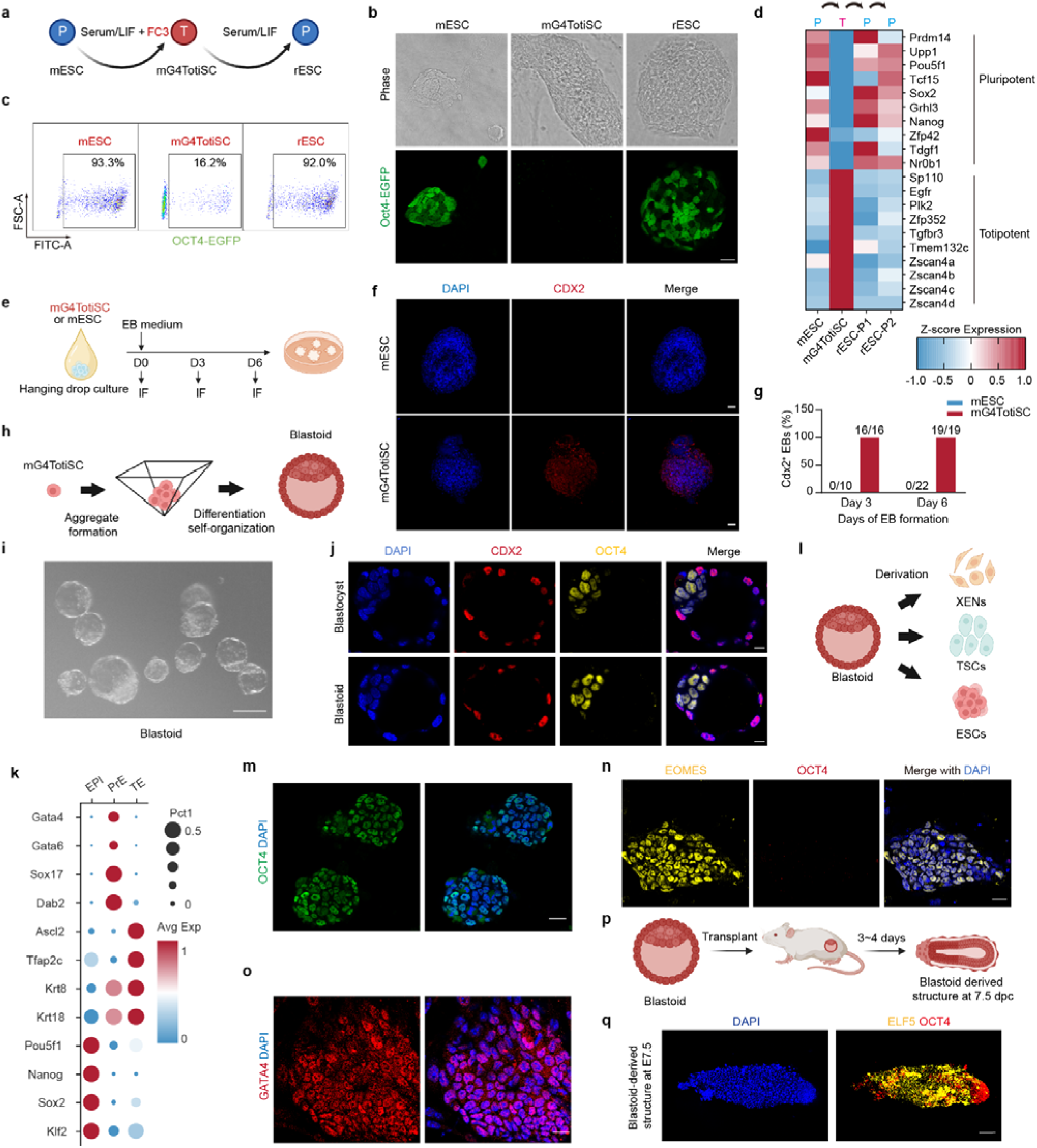
Differentiation and self-organization ability of mG4TotiSCs. **a,** Schematic of the reversible system used to investigate the totipotent/pluripotent transition. rESCs, reversed ESCs; P, pluripotent; T, totipotent. **b**, Morphology of OCT4-EGFP mESCs during the transition from mESCs (P0) to mG4TotiSCs (P6) and their conversion back to rESCs (rP2). Green color, EGFP. Scale bar, 25 μm. **c**, FACS analysis of OCT4-EGFP mESCs during the transition from mESCs (P0) to mG4TotiSCs (P6) and their conversion back to rESCs (rP2). **d**, Heatmap showing the relative expression of representative pluripotent and totipotent genes in mouse mESCs (P0), mG4TotiSCs (P6) and rESCs (rP1, rP2). **e,** Schematic diagram showing the direct differentiation and identification of **embryoid bodies (EBs)** from mESCs and mG4TotiSCs. IF, immunofluorescence. **f,** Immunofluorescence imaging of CDX2 expression in EBs derived from mG4TotiSCs or mESCs. Red color, CDX2. Scale bar, 200 μm. **g,** Percentage of CDX2^+^ EBs on days 3 and 6. **h**, Schematic diagram showing the differentiation and self-organization of mG4TotiSCs to derive blastoids. **i**, Morphology of mG4TotiSC-derived blastoids. Scale bar, 100 μm. **j**, Immunofluorescence imaging of CDX2 and OCT4 expression in blastocysts or blastoids. Scale bar, 10 μm. **k**, Bubble plot from scRNA-seq showing the relative expression of representative EPI, PrE and TE genes in the mG4TotiSC-derived blastoids. **l**, Schematic diagram showing the lineage-specific differentiation of mG4TotiSC-derived blastoids. **m**, Immunofluorescence imaging of OCT4 expression in ESCs differentiating from mG4TotiSC-derived blastoids. Scale bar, 25 μm. **n**, Immunofluorescence imaging of EOMES and OCT4 expression in TSCs differentiating from mG4TotiSC-derived blastoids. Scale bar, 25 μm. **o**, Immunofluorescence imaging of GATA4 expression in XEN cells differentiating from mG4TotiSC-derived blastoids. Scale bar, 25 μm. **p**, Schematic diagram showing transplantation of mG4TotiSC-derived blastoids into the uteri of pseudopregnant mice and its post-implantation development. **q**, Immunofluorescence imaging of ELF5 and OCT4 expression in blastoid-derived structure at 7.5 dpc. Scale bar, 100 μm.

A key functional characteristic of TotiSCs is their capacity to generate both embryonic and extraembryonic lineages, including trophoblast stem cells (TSCs), a capability absent in conventional pluripotent stem cells. To evaluate this potential in mG4TotiSCs, we subjected them to established TSC differentiation conditions (**Extended Data Fig. 3d**). Time-course RT-qPCR analysis revealed upregulation of canonical trophoblast lineage determinants, including Cdx2, Elf5, Tfap2c, and Esx1, in differentiating mG4TotiSCs.In contrast, parallel cultures of mESCs under the same conditions showed only minimal induction of these markers (**Extended Data Fig. 3e**). Furthermore, we conducted embryoid body (EB) formation assays under standard suspension culture conditions (**Fig. 3e**). Strikingly, while CDX2 protein expression was nearly undetectable in EBs derived from mESCs, alomost all mG4TotiSC-derived EBs demonstrated strong CDX2 expression **(Fig. 3f-g**). Collectively, these findings provide compelling evidenve that mG4TotiSCs possess a bipotential differentiation capacity in vitro, giving rise to both pluripotent derivatives and extraembryonic trophoblast lineages.

### mG4TotiSCs generate blastoids autonomously without external signalling

A hallmark of TotiSCs is their capacity to self-organize into blastocyst-like structures, termed blastoids.The blastoids faithfully recapitulate the architecture, lineage composition, and developmental potential of natural blastocysts. We systematically evaluated the blastoid-forming competence of mTotiSCs using a microwell-based aggregation assay: single-cell mTotiSCs were seeded into microwells at densities exceeding five cells per well to promote controlled formation of compact multicellular aggregates **(Fig. 3h**). Blastoid induction was carried out in a well-verified medium adapted from previously established protocols^40^. Morphological assessment revealed blastoids began to form on the third day after seeding, and fully cavitated spherical blastocyst-like structures emerging between days 5 and 6 **(Fig. 3i**), measuring 100–120 μm in diameter. Approximately 20% of seeded aggregates progressed to morphologically mature blastoids. Immunofluorescence analysis confirmed spatially resolved lineage specification: CDX2 was robustly expressed in the outer trophectoderm (TE)-like layer, while OCT4 localized specifically to the ICM-like compartment, mirroring the canonical spatial patterning of natural blastocysts **(Fig. 3j**). scRNA-seq of mG4TotiSC-derived blastoids further validated their molecular fidelity, revealing transcriptionally distinct populations corresponding to epiblast (EPI), primitive endoderm (PrE), and TE lineages **(Fig.3k; Extended Fig. 3f**). Critically, when dissociated and replated under lineage-specific differentiation conditions **(Fig. 3l**), individual blastoids gave rise to stable, self-renewing ESCs **(Fig. 3m; Extended Data Figure 3g**), TSCs **(Fig. 3n; Extended Data Figure 3h**), and extraembryonic endoderm (XEN) cells **(Fig. 3o; Extended Data Figure 3i**). Moreover, after transplantation into the uteri of pseudopregnant mice **(Fig. 3p**), mG4TotiSC-derived blastoids exhibited efficient uterine implantation and initiated development, generating embryo-like structures closely resembling E7.5 conceptuses in morphology and tissue organization **(Fig. 3q; Extended Fig. 3j-k**). Collectively, these findings establish that mG4TotiSCs possess intrinsic self-organizing capacity and functional totipotency, by clarifying their ability to generate both embryonic and extraembryonic lineages in vitro and support peri-implantation embryogenesis in vivo.

### mG4TotiSCs contribute to both embryonic and extraembryonic lineages in chimeric mice

The most stringent functional criterion for assessing totipotency in stem cells is their capacity to contribute to embryonic and extraembryonic lineages in chimeric embryos^41^.. To assess whether mG4TotiSCs meets the criterion, we performed an in vitro chimeric blastocyst assay. Specifically, we microinjected either one or three EGFP-labeled mG4TotiSCs, or control mESCs, into 8-cell-stage embryos derived from ICR mice (**Fig. 4a**). The chimeric embryos were then cultured in vitro for 36-48 hours to allow lineage integration and differentiation prior to analysis. Immunofluorescence staining for the TE-specific marker CDX2 revealed that 92% of blastocysts injected with a single EGFP□ mG4TotiSC contained EGFP□ cells co-expressing CDX2 in the TE layer, along with EGFP□ cells localized to the OCT4^+^ ICM compartments (**Fig. 4b, c**).This result demonstrating unequivocal contribution to both extraembryonic and embryonic lineages. In stark contrast, EGFP□ mESCs exclusively integrated into the ICM and consistently failed to colonize or differentiate into the CDX2□ TE compartment under identical experimental conditions. Comparable lineage-restricted integration was observed when three mESCs were injected, whereas multi-cell mG4TotiSC injections yielded proportionally higher chimerism without compromising TE/ICM bifurcation (**Extended Data Fig. 4a, b**). These results provide direct evidence that mG4TotiSCs possess true totipotent potential, as defined by their ability to generate derivatives of both embryonic and extraembryonic cells within a developing blastocyst context.

**Fig. 4.**
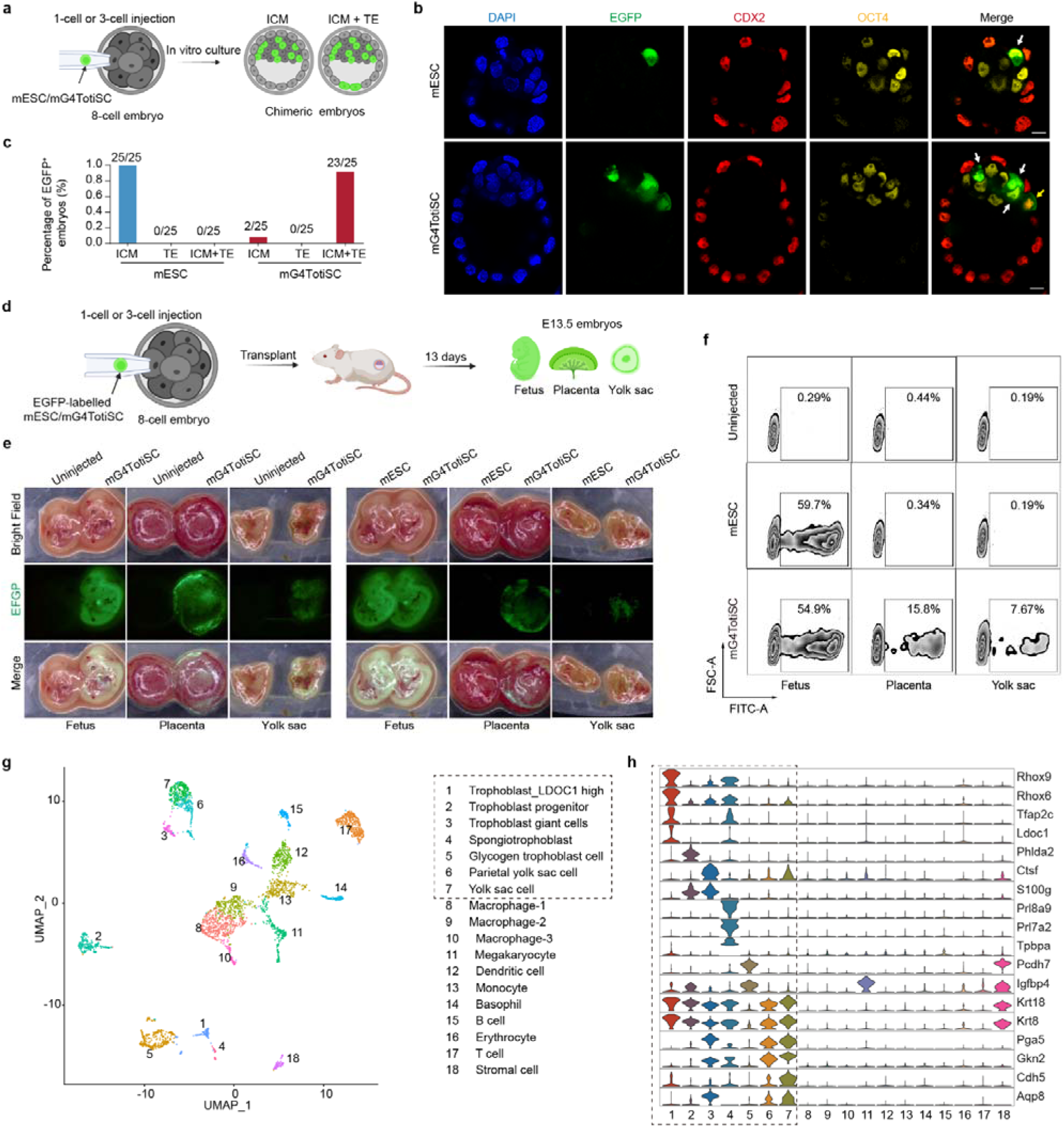
mG4TotiSCs have bidirectional differentiation potential in chimeric embryos. **a,** Schematic diagram of the experimental design used to generate chimeric blastocysts with EGFP^+^ mG4TotiSCs or mESCs. **b,** Representative immunofluorescence images showing the expression of CDX2 and OCT4 in chimeric blastocysts generated from 8-cell embryos injected with a single EGFP^+^ mG4TotiSC or mESC in vitro. Green color, EGFP. Red color, CDX2. Yellow color, OCT4. Scale bars, 10 μm. **c,** Fraction of chimeric blastocysts in which a single injected EGFP^+^ mG4TotiSC or mESC contributed to the ICM, TE or both. The fraction is indicated above each bar. **d,** Schematic diagram of the experimental design used to generate chimeric E13.5 embryos with EGFP^+^ mG4TotiSCs or mESCs. **e,** Images of chimeric conceptuses derived from 8-cell embryos injected with EGFP^+^ mESCs or mG4TotiSCs with an uninjected conceptus included for reference. **f,** FACS analysis of EGFP^+^ cells in chimeric conceptuses. The percentages of EGFP^+^ cells derived from donor mESCs and mG4TotiSCs are shown, and cells isolated from uninjected concepti served as a control. **g,** Uniform manifold approximation and projection (UMAP) plot showing the 18 main clusters. The dotted line indicates extraembryonic cell lineages. **h,** Violin plots displaying the relative expression distributions of specific marker genes in clusters of extraembryonic lineages. The dotted line highlights the extraembryonic cell lineages.

Next, we assessed the bidirectional developmental potential of mG4TotiSCs in post-implantation chimeric embryos in vivo. During post-implantation development, the TE lineage gives rise to extraembryonic tissues, including the placenta, whereas the ICM differentiates into both the embryonic fetus and the extraembryonic yolk sac. To evaluate lineage contribution beyond the blastocyst stage, we transferred 8-cell-stage chimeric embryos, generated by microinjecting one or three EGFP-labeled mG4TotiSCs or control mESCs, into the uteri of pseudopregnant mice. Embryos were harvested at embryonic day 13.5 (E13.5) for comprehensive tissue-level analysis (**Fig. 4d**). Consistent with totipotent capacity, mG4TotiSC-derived EGFP□ cells were robustly detected across all major compartments: both the embryonic fetus and the extraembryonic placenta and yolk sac (**Fig. 4e; Extended Data Fig. 4c**). In contrast, mESC-derived EGFP□ cells were exclusively restricted to the fetal compartment, with no detectable contribution to extraembryonic tissues. Quantitative flow cytometric analysis further revealed that EGFP□ cells constituted 54.9% of fetal cells, 15.8% of placental cells, and 7.67% of yolk sac cells in mG4TotiSC chimeras. However, in mESC chimeras, EGFP□ cells accounted for 59.7% of fetal cells and were absent in extraembryonic tissues (**Fig. 4f**). To validate extraembryonic differentiation at the cellular level, we performed immunofluorescence staining for proliferin, a well-established marker of trophoblast giant cells, on cryosections of E13.5 placentas. EGFP□ mG4TotiSC derivatives were abundantly present in both the labyrinthine and junctional zones of the placenta, where they co-expressed proliferin, thereby confirming functional integration into terminally differentiated trophoblast lineages (**Extended Data Fig. 4d**. In contrast, no mESC-derived EGFP^+^ cells were found in the placenta. Notably, mG4TotiSCs also significantly contributed to chimeric fetal organs, including brain, heart, and liver, as well as to the germline, enabling the generation of viable adult offsprings (**Extended Data Fig. 4e–g**). Collectively, these in vivo chimerism assays demonstrate that mG4TotiSCs possess comprehensive developmental competence, making functional contributions to embryonic, extraembryonic, and germline lineages throughout gestation.

Finally, to comprehensively resolve the cellular diversity and lineage trajectories of mG4TotiSC-derived extraembryonic progeny at single-cell resolution, we performed scRNA-seq on EGFP□ cells from E13.5 chimeric placentas and yolk sacs isolated via fluorescence-activated cell sorting (FACS). Unsupervised graph-based clustering coupled with rigorous annotation using canonical lineage-defining markers revealed seven transcriptionally distinct mG4TotiSC-derived extraembryonic cell types (**Fig. 4g, h; Extended Data Fig. 5**): (i) LDOC1-high trophoblast cells, co-expressing epithelial keratins (KRT8, KRT18), Rhox family transcription factors (Rhox6, Rhox9), and elevated LDOC1; (ii) trophoblast progenitor cells, marked by Tfap2c and Phlda2; (iii) trophoblast giant cells, defined by Ctsf expression; (iv) spongiotrophoblast cells, characterized by Tpbpa, Prl8a9, and Prl7a2; (v) glycogen trophoblast cells, identified by Pcdh7 and Igfbp4; (vi) parietal yolk sac cells, expressing Cdh5 and Gkn2; and (vii) yolk sac cells, distinguished by Cdh5, Gkn2, and Phldb2. Notably, we also detected low-abundance EGFP□ cells of embryonic origin, such as macrophages, dendritic cells, stromal cells and erythrocytes, indicating contribution to embryonic-extraembryonic interface tissues. Collectively, these high-resolution transcriptomic data demonstrate that mG4TotiSCs generate a full spectrum of extraembryonic lineages in vivo, with molecular fidelity to their native counterparts, and robustly contribute to embryonic tissues as well. This comprehensive developmental competence fulfills the most stringent functional criteria for totipotency, thereby establishing mG4TotiSCs as bona fide TotiSCs.

### The induction of mG4TotiSCs from mESCs is mediated by FC3-induced Gata2 mRNA unwinding

After confirming that FC3 induces the functional transition from mESCs to mG4TotiSCs, we investigated the underlying molecular mechanism. Our previous research showed that FC3 enhances the translational efficiency of mRNAs containing rG4 structures^34^. To systematically quantify genome-wide translational changes associated with this transition, we conducted an active ribosome profiling assay (RiboLace), a high-resolution sequencing method for quantifying mRNA translation levels, on matched mG4TotiSC and mESC cultures. Consistent with FC3’s rG4-targeting activity, we found that transcripts with predicted rG4 motifs in their 5′-UTRs, CDS, or 3′-UTRs had significantly higher translation efficiency in mG4TotiSCs relative to mESCs (**Extended Data Fig. 6a**). Notably, among CDS-embedded rG4-containing genes, 137 showed increased translation efficiency, whereas only 31 displayed reduced translation efficiency, indicating a strong directional bias toward translational activation. These data suggest that FC3-mediated enhancement of rG4-driven translation is a central regulatory node in the pluripotency-to-totipotency transition.

We further compared translational efficiency profiles between early 2-cell blastomeres and the ICM to assess developmental stage–specific regulation by rG4s. Using a stringent threshold, fold change ≥ 1.70 for two-cell–enriched translation and ≤ 0.588 for ICM-enriched translation, we defined two non-overlapping gene sets: a 2-cell–preferentially translated set and an ICM–preferentially translated set. Strikingly, 16% of genes in the two-cell–preferentially translated set harbored rG4 motifs in their transcript sequences (**Fig. 5a**), whereas only 13% of genes in the ICM–preferentially translated set contained such motifs, a proportion statistically indistinguishable from the genome-wide background frequency (∼13%). This enrichment specifically in the two-cell–translated cohort suggests that rG4 structures are preferentially associated with transcripts subject to dynamic translational activation during the two-cell stage, rather than passive retention or general abundance. Consequently, FC3-mediated rG4 unwinding likely facilitates sustained translation of these developmentally critical transcripts, thereby stabilizing the 2-cell–like molecular state in mG4TotiSCs.

**Fig. 5.**
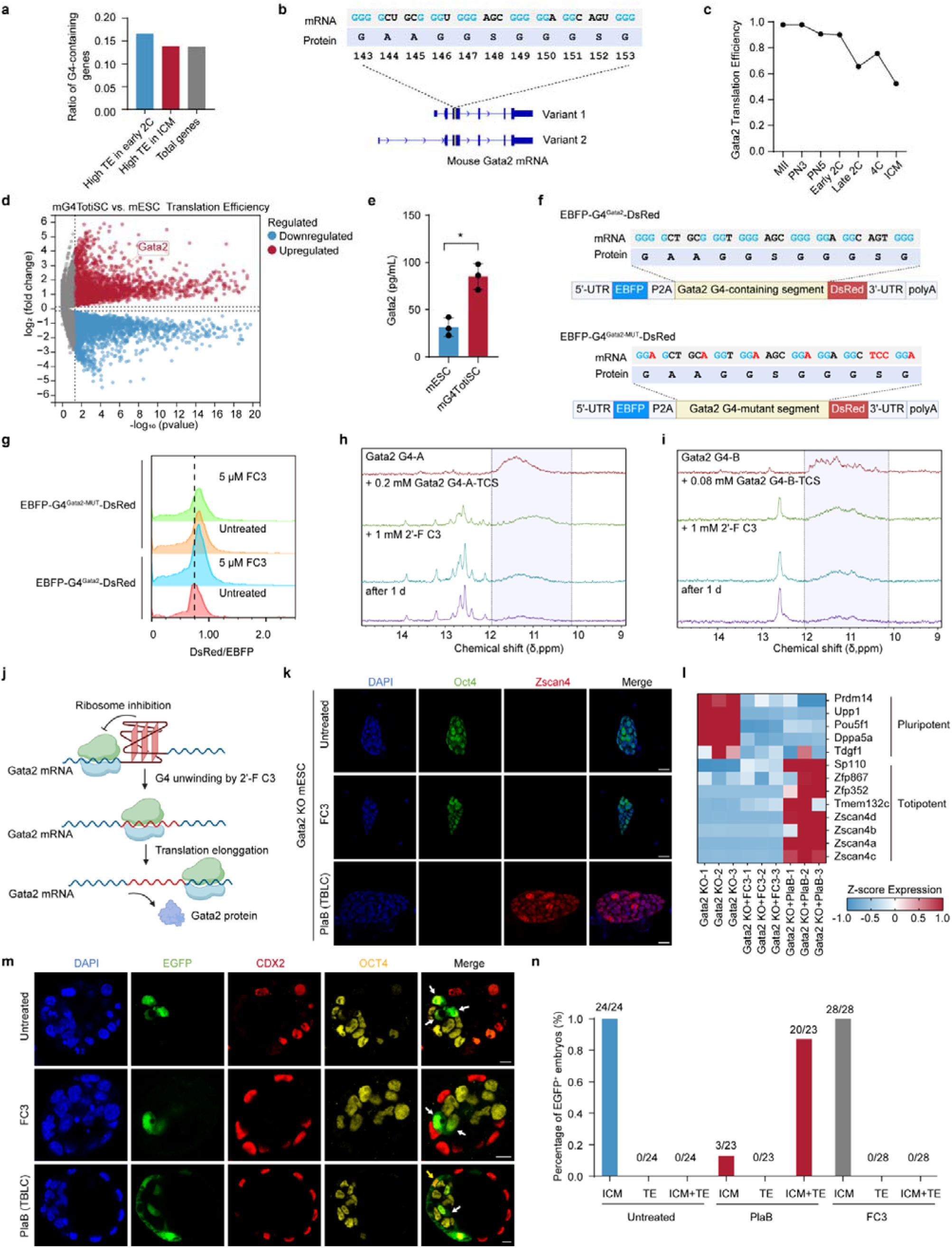
FC3-induced totipotency transition of mG4TotiSCs is GATA2-rG4-dependent. **a**, Ratio of G4-containing genes in genes with high translational efficiency in mouse early 2-cell embryos and ICMs. **b**, Gata2 G4 motif and the corresponding amino acids. **c**, GATA2 translation efficiency during early embryo development in mice. All the data are from a published resource^43^. **d**, RiboLace analysis of mG4TotiSCs and mESCs and volcano plot displaying the global changes in mRNA translation efficiency. Red and blue dots indicate upregulated and downregulated genes with p < 0.05. **e**, ELISA analysis of GATA2 protein expression in mG4TotiSCs and mESCs. The Gapdh internal control protein was analysed via the same blot. **f**, Schematic diagram of the reporter mRNA design used to evaluate the unfolding of Gata2 G4 structures and the upregulation of translation. **g**, Histogram of the ratio of the dsRed fluorescence intensity to the EBFP fluorescence intensity detected via FACS. After reporter mRNA transfection, HEK293FT cells were treated with 5 μM FC3 for 12 h. **h**, NMR spectra of Gata2 G4-A oligos sequentially incubated with truncated complementary strands (TCSs) and FC3. **i**, NMR spectra of Gata2 G4-B oligos sequentially incubated with TCSs and FC3. **j**, Schematic diagram of the mechanism of the FC3-induced increase in Gata2 translation. **k**, Immunofluorescence imaging of Oct4 and Zscan4 in Gata2 KO mESCs and Gata2 KO mESCs treated with FC3 or PlaB. Scale bar, 20 μm. **l**, Changes in the transcription of pluripotency and totipotency genes among Gata2 KO mESCs and Gata2 KO mESCs treated with FC3 or PlaB. **m**, Representative immunofluorescence images showing the expression of CDX2 and OCT4 in chimeric blastocysts generated from 8-cell embryos injected with multiple EGFP^+^ Gata2 KO mESCs or Gata2 KO mESCs treated with FC3 or PlaB. Scale bar, 10 μm. **n**, Fractions of chimeric blastocysts in which EGFP^+^ Gata2 KO mESCs or Gata2 KO mESCs treated with FC3 or PlaB contributed to the ICM, TE or both. The fraction is indicated above each bar.

Several functional proteins such as GATA2^39^, DUX^12^, EIF3H^42^, DPPA2/4^9^, NELFA^11^, and ATR^42^, have previously been linked to the induction of totipotency. To pinpoint the essential regulators responsible for the emergence of mG4TotiSCs, we focused on these functionally validated candidates. Using pqsfinder^27^ with a G-score threshold of 52, we systematically scanned the transcript sequences of these candidate totipotency-associated genes for rG4 motifs. Notably, only Gata2 mRNA contained a high-confidence rG4 motif, located within its CDS and conserved across two splice isoforms (**Fig. 5b**). Strikingly, this motif is largely built from consecutive GGG glycine codons, even though GGC (47%) is the most frequently used glycine codon in Gata2, whereas GGG accounts for only 21%, suggesting evolutionary selection to enrich GGG to stabilize the rG4 structure. Consistent with a regulatory role, translational efficiency of Gata2 mRNA gradually declines during early embryogenesis^43^: it reaches its highest level from the zygote to two-cell stage and falls to approximately half by the ICM stage (**Fig. 5c**), paralleling the developmental narrowing of totipotency To confirm structural integrity, we synthesized two oligonucleotides spanning the predicted Gata2 rG4 sequence and performed ¹H NMR spectroscopy. Both oligos displayed typical G4 signals between 10–12 ppm, confirming intramolecular G-quadruplex formation. These signals disappeared entirely upon G-to-A mutagenesis, confirming sequence-specific G4 folding **(Extended Data Fig. 6b, c**). Furthermore, single-molecule rolling circle amplification (RCA)^44^ imaging revealed basal, heterogeneous Gata2 mRNA expression in mESCs (**Extended Data Fig. 6d**), Earlier work has also shown that inhibition of miR-34a, a direct post-transcriptional repressor of Gata2, promotes the acquisition of totipotent-like features^39^. Taken together, these orthogonal lines of evidence, bioinformatic prediction, structural validation, developmental translation dynamics, and functional genetic modulation, converge to establish the Gata2 rG4 motif as a cis-regulatory element governing translational control of a key totipotency gatekeeper.

Next, we investigated the mechanistic role of FC3 in regulating Gata2 expression during mG4TotiSC induction. RiboLace profiling revealed a significant increase in translational efficiency of Gata2 mRNA in mG4TotiSCs relative to mESCs (**Fig. 5d**). Consistently, quantitative ELISA and immunoblotting confirmed a strong elevation in GATA2 protein abundance (**Fig. 5e; Extended Data Fig. 6e**). These findings consolidate the role of GATA2 in facilitating the shift from pluripotency to totipotency and refine our understanding of its basal, yet dynamically regulated, expression in mESCs. To directly examine FC3’s ability to regulate translation via rG4, we designed a dual-fluorescent reporter construct in which EBFP is fused upstream of a Gata2-derived CDS-embedded rG4 motif, followed by DsRed (**Fig. 5f**). In this design, the rG4 structure acts as a cis-repressive element: its stable folding sterically impedes ribosome progression, thereby suppressing downstream DsRed synthesis. Upon FC3-mediated rG4 resolution, ribosomal transit is restored, boosting DsRed translation and elevating the DsRed/EBFP fluorescence intensity ratio, a quantitative proxy for translational derepression. As predicted, treating mESCs with FC3 for 12 hours led to an approximately 20% increase in the DsRed/EBFP ratio in mESCs (**Fig. 5g; Extended Data Fig. 6f**). Critically, mutation of the core G-tracts within the rG4 motif abolished both basal repression and FC3 responsiveness, confirming sequence-specificity. Furthermore, ¹H NMR unwinding assays demonstrated that FC3 facilitated the unwinding of Gata2 rG4 structures, which was direct biophysical evidence of FC3-induced structural destabilization of Gata2 rG4 (**Fig. 5h-i**). Collectively, this combination biochemical, genetic, and biophysical evidence demonstrates that FC3 promotes GATA2 protein synthesis by selectively resolving the inhibitory rG4 structure in the Gata2 coding sequence, thereby enabling efficient ribosome elongation (**Fig. 5j**).

As established in prior research, GATA2 serves as a master transcriptional regulator of MERVL retrotransposon-derived transcripts and acts as a critical upstream activator within totipotency gene network^39^. To assess whether GATA2 is functionally required for FC3-induced reprogramming into mG4TotiSCs, we performed CUT&Tag analysis to map GATA2 chromatin binding in mG4TotiSCs compared with control mESCs. Strikingly, GATA2 exhibited robust and specific enrichment at the MERVL-int and MT2_Mm transposon elements, in mG4TotiSCs (**Extended Data Fig. 7a-c**), whereas no significant binding was detected at these sites in mESCs. To rigorously test GATA2 dependence, we generated Gata2 knockout mESCs (Gata2 KO mESCs) (**Extended Data Fig. 6d**) and subjected them to FC3-based reprogramming under SLFM conditions. As a mechanistic control, we also applied the PlaB protocol, which has been previously demonstated to generate TBLCs^4^, to the same Gata2 KO line. Immunofluorescence analysis of ZSCAN4, a well-validated marker of totipotency, revealed that FC3 treatment failed to induce ZSCAN4 expression in Gata2 KO mESCs, whereas PlaB treatment strongly activated ZSCAN4 even in the absence of GATA2 (**Fig. 5k**). This clear contrast demonstrates that FC3-driven totipotency acquisition strictly depends on GATA2, whereas PlaB-mediated reprogramming bypasses this requirement. Bulk RNA-seq further supported this distinction: FC3-treated Gata2 KO mESCs displayed gradual downregulation of core pluripotency genes but completely failed to activate key totipotency transcripts such as Zscan4 cluster members, and MERVL-derived RNAs, suggesting a block at the transcriptional initiation step of the totipotency program (**Fig. 5i; Extended Data Fig. 7e-f**). In contrast, PlaB treatment robustly upregulated totipotency-associated genes and MERVL-derived transcripts while markedly suppressing canonical pluripotency factors. To assess functional developmental potential, we performed chimeric blastocyst assays by microinjecting three treated cells into ICR 8-cell embryos and culturing them to the blastocyst stage. Consistent with the molecular and cellular phenotypes, FC3-treated Gata2 KO mESCs completely failed to integrate into or differentiate within the TE lineage, with no EGFP□ TE cells detected in any chimeric blastocyst (**Fig. 5m-n**). Conversely, PlaB-treated Gata2 KO mESCs efficiently contributed to both the inner cell mass (ICM) and TE compartments, with ∼90% of chimeric blastocysts showing dual-lineage contribution (**Fig. 5m-n**). These functional experiments provide definitive evidence that GATA2 is not universally required for totipotency, but rather serves as a pathway-specific effector essential for FC3-driven reprogramming. Collectively, our integrated genetic, transcriptional, and developmental analyses establish the Gata2 rG4 motif as a critical node through which FC3 selectively licenses the totipotent transition.

### G4 unwinding induces human totipotency

Currently, the robust generation and sustained cultivation of human TotiSCs, capable of generating both embryonic and extraembryonic lineages in vivo, remains difficult. To date, only two types of totipotent-like cells have been described: 8-cell-like cells (8CLCs) and human totipotent blastomere-like cells (hTBLCs). To test whether unwinding of rG4s could stably reprogramme human cells toward a totipotent state, we performed a systematic screen of FC3-based culture conditions. We identified a chemically defined, feeder-free culture system, E8 medium supplemented with 5 μM FC3, 2 μM Y-27632, 2 μM minocycline, and 1 μM CHIR99021, that enables efficient, reproducible conversion of conventional hESCs into hG4TotiSCs within 7 days on Matrigel-coated plates. Immunofluorescence imaging confirmed the near-complete loss of core pluripotency transcription factors OCT4, alongside pronounced nuclear accumulation of the ZGA markers ZBTB16 in hG4TotiSCs (**Fig. 6a**). After multiple passages, hG4TotiSC exhibited no obvious apoptosis and maintained a normal diploid karyotype (**Extended Data Fig. 8a-b**). Transcriptome-wide analysis revealed comprehensive transcriptional rewiring: hG4TotiSCs strongly upregulated pre-ZGA and ZGA-associated genes while suppressing canonical pluripotency network components (**Fig. 6b; Extended Data Fig. 8c**), which was further validated by scRNA-seq (**Fig. 6c; Extended Data Fig. 8d**). At the epigenetic level, as anticipated, regulatory regions of core totipotency-associated genes (PLAGL1, PRTG, RAI1, and ZBTB16) exhibited significantly increased accessibility in mG4TotiSCs, whereas loci associated with key pluripotency factors (EPCAM, NANOG, POU5F1, and SOX2) showed pronounced reductions in accessibility (**Extended Data Fig. 8e**). In all, these results provide molecular and epigenetic evidence for the totipotent identity of hG4TotiSCs.

**Fig. 6.**
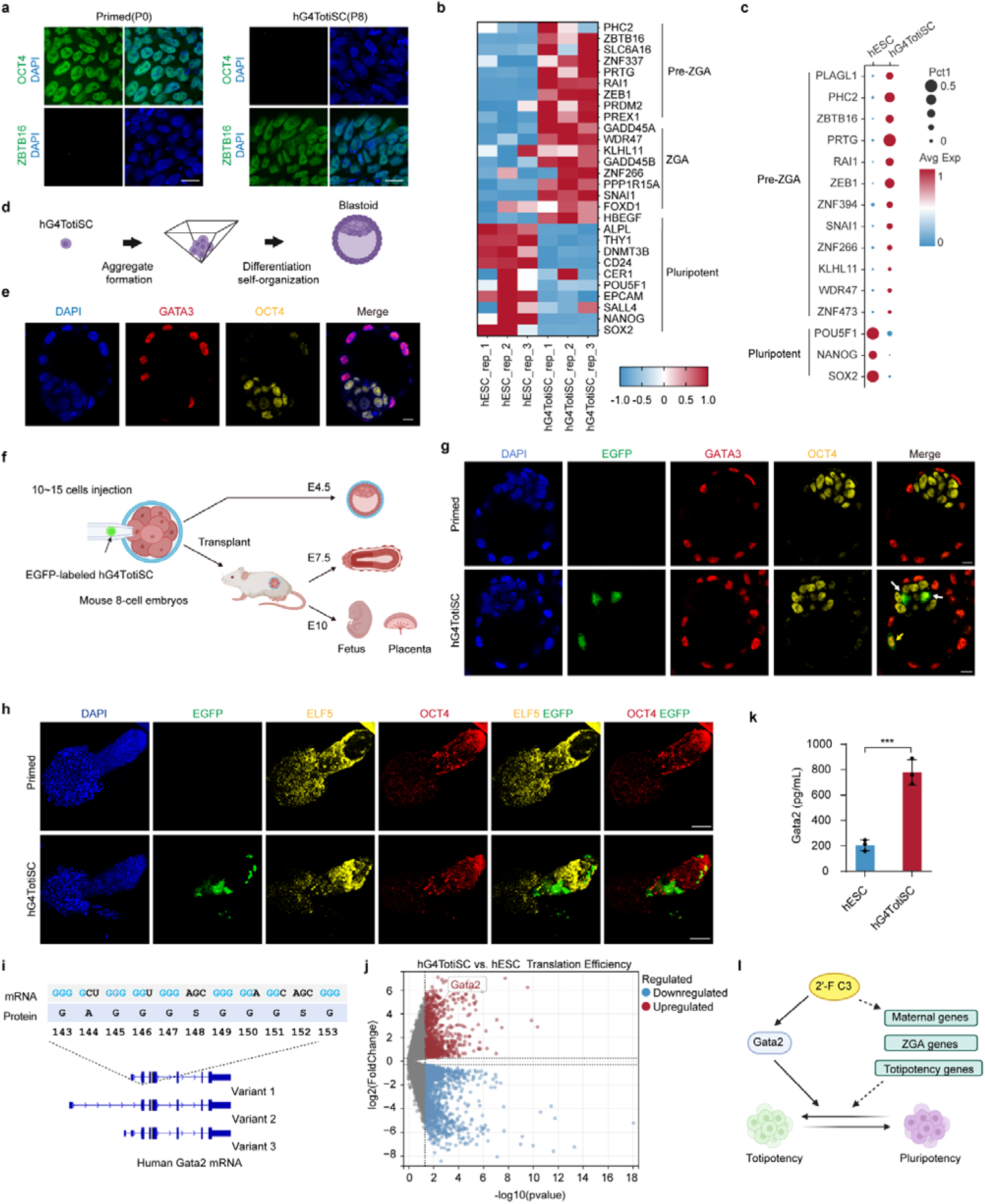
FC3 induces human totipotency. **a**, Representative immunofluorescence images of OCT4 and ZBTB16 in hESCs or hG4TotiSCs. Scale bar, 25 μm. **b**, Heatmap showing the relative transcription levels of representative Pre-ZGA, ZGA and pluripotent genes in hESCs and hG4TotiSCs. **c**, Bubble plot from scRNA-seq showing the relative expression of representative pre-ZGA and pluripotent genes in the hESC and hG4TotiSC clusters. **d**, Schematic diagram showing the differentiation and self-organization of hG4TotiSCs to derive blastoids. **e**, Immunofluorescence imaging of GATA3 and OCT4 expression in hG4TotiSC-derived blastoids. Scale bar, 10 μm. **f**, Schematic diagram of the experimental design used to generate chimeric E4.5, E7.5 and E13.5 embryos with EGFP-labeled hG4TotiSCs. **g,** Representative immunofluorescence images showing the expression of GATA3 and OCT4 in chimeric blastocysts generated from 8-cell embryos injected with EGFP-labeled hG4TotiSCs or hESCs in vitro. Green color, EGFP. Red color, GATA3. Yellow color, OCT4. Scale bars, 10 μm. **h**, Immunofluorescence imaging of ELF5 and OCT4 expression in E7.5 chimeras generated from 8-cell embryos injected with EGFP-labeled hG4TotiSCs or hESCs. Scale bar, 100 μm. **i**, Human GATA2 G4 motif and the corresponding amino acids. **j**, RiboLace analysis of hG4TotiSCs and hESCs and volcano plot displaying the global changes in mRNA translation efficiency. Red and blue dots indicate upregulated and downregulated genes with p < 0.05. **k**, ELISA analysis of GATA2 protein expression in hG4TotiSCs and hESCs. **l**, Model of the role of FC3 in reprogramming cell fate to a totipotent state.

We next assessed the spontaneous lineage differentiation potential of hG4TotiSCs. When cultured on Matrigel in E8 medium without additional components, hG4TotiSCs underwent rapid and synchronous multilineage differentiation within 3–4 days (**Extended Data Fig. 9a**), generating cells that expressed lineage-specific markers of the EPI (OCT4□), TE (GATA3□), and PrE (GATA6□), as confirmed by immunofluorescence imaging (**Extended Data Fig. 9b**). This tri-lineage differentiation potential was further validated at single-cell resolution (**Extended Data Fig. 9c-d**): scRNA-seq analysis identified distinct transcriptional clusters corresponding to EPI-, TE-, and PrE-like states. To evaluate self-organization ability, we examined whether hG4TotiSCs could form blastoids. Following gentle dissociation, 20–25 hG4TotiSCs were seeded per microwell to promote controlled aggregate formation (**Fig. 6d**). When cultured in G-2 Plus medium, ∼10% of aggregates developed into polarized, cavity-containing blastoids by day 5–6 (**Extended Data Fig. 9e**). Immunostaining confirmed spatially restricted expression of lineage markers (**Fig. 6e**): GATA3□ cells organized into a cohesive outer layer resembling TE, while an inner OCT4□ cluster recapitulated ICM architecture, establishing both structural polarity and molecular fidelity to native human blastocysts. Finally, we rigorously evaluated developmental potency of hG4TotiSCs using xenogeneic chimerism assays. 10 to 15 EGFP-labeled hG4TotiSCs were microinjected into ICR mouse 8-cell embryos (**Fig. 6f**). In vitro, chimeric blastocysts (E4.5) exhibited widespread EGFP□ contribution to both OCT4□ (ICM/EPI) and GATA3□ (TE) compartments (**Fig. 6g**), unlike control hESCs, which failed to integrate or survive beyond 48 hours. Upon embryo transfer into pseudopregnant recipients, hG4TotiSC-derived chimeras progressed to E7.5 (**Fig. 6h**), where EGFP□ cells co-expressed OCT4 and ELF5 in nascent embryonic and extraembryonic ectoderm domains. Critically, at E10.5, EGFP□ cells contributed robustly to both embryonic tissues (**Extended Data Fig. 9f-h**), including extraembryonic lineages such as KRT7□ trophoblast giant cells in the placenta, as well as PAX6□ neuroectoderm and GATA4□ endodermal lineages. Collectively, these functional assays demonstrate that hG4TotiSCs possess bona fide totipotent developmental potential, capable of generating not only embryonic lineages but also functionally integrated extraembryonic tissues in vivo.

Mechanistically, we identified a conserved rG4 motif within the CDS of human GATA2 mRNA (**Fig. 6i**), located in the coding region of amino acids 143-153, just like in mice. Consistently, RiboLace profiling demonstrated a significant enhancement in GATA2 translational efficiency, confirming post-transcriptional upregulation (**Fig. 6j**). Quantitative ELISA revealed a ∼3.2-fold increase in GATA2 protein abundance in hG4TotiSCs relative to parental hESCs (**Fig. 6k**). Crucially, ¹H NMR confirmed that the predicted GATA2 rG4 sequence adopts a stable, intramolecular G4 conformation (**Extended Data Fig. 9i-j**), and that FC3 directly destabilizes this structure, as shown by loss of characteristic G4 signals (10–12 ppm) (**Extended Data Fig. 9k-l**). Together, these orthogonal data, spanning bioinformatic prediction, quantitative assay, translational profiling, and biophysical validation, establish that FC3-mediated resolution of the GATA2 rG4 motif drives enhanced GATA2 synthesis, thereby enabling totipotency acquisition. Critically, parallel analyses in mouse mG4TotiSCs revealed an identical dependency on Gata2 rG4 unwinding for functional reprogramming, demonstrating evolutionary conservation of this rG4–GATA2–totipotency axis across mammalian species.

## Discussion

Owing to limited mechanistic understanding of the molecular determinants of stem cell totipotency, the robust and scalable derivation of expandable totipotent stem cells (TotiSCs) from somatic or pluripotent precursors remains a longstanding challenge. Given accumulating evidence that rG4s function as dynamic regulators in stem cell fate decisions, including lineage priming and differentiation, we hypothesized that chemical modulation of rG4 stability could offer a viable strategy to unlock totipotency. In this study, we identify rG4 unwinding as a previously unrecognized, evolutionarily conserved regulatory node governing totipotency acquisition in both mouse and human systems. By supplementing conventional mESC or hESC culture media with the selective rG4-unwinding small molecule FC3, we achieved highly efficient and reproducible reprogramming into stable, self-renewing totipotent stem cells, designated mG4TotiSCs and hG4TotiSCs, respectively. Both lines exhibited long-term genomic stability: after ≥10 consecutive passages under standard conditions, they maintained normal diploid karyotypes and displayed apoptosis rates indistinguishable from parental controls. Transcriptomic and epigenomic profiling confirmed that mG4TotiSCs and hG4TotiSCs closely recapitulate the molecular signature of in vivo 2-cell blastomeres. Functional validation using blastoid formation and chimerism assays demonstrated that both mG4TotiSCs and hG4TotiSCs contribute robustly to embryonic and extraembryonic lineages, fulfilling the gold-standard criterion for functional totipotency. Mechanistically, FC3 directly resolves unfold evolutionarily conserved rG4 structures within the CDS of GATA2 mRNA, enhancing ribosome loading and translational to elevated GATA2 protein levels, which are essential for totipotency induction. Critically, genetic ablation of Gata2 abolished FC3-mediated reprogramming in mESCs, whereas PlaB-induced totipotency remained fully intact, underscoring pathway specificity. Collectively, our work provides the first rG4-targeted chemical protocol for generating expandable, functionally validated TotiSCs from non-germline sources, and identifies rG4-mediated translational control of GATA2 as a fundamental axis regulating the pluripotency–totipotency transition across mammals.

Optimizing the derivation of TotiSCs from non-germline sources in vitro represents a pivotal advance for developmental biology and regenerative medicine. To date, only a limited number of chemical protocols, including D-EPSC, L-EPSC, TPS, TLSC, CiTotiSC, h8CLC, mTBLC, and hTBLC, have demonstrated capacity to induce totipotent-like states. Notably, these rely on empirically assembled multi-target cocktails that modulate distinct signaling axes, such as retinoic acid receptor activation (TTNPB), Wnt/β-catenin potentiation (1-azakenpaullone), and NF-κB pathway suppression (WS63) in CiTotiSC generation^3^. This polypharmacological complexity compromises reproducibility, obscures mechanistic interpretation, and hinders translational scalability. In contrast, our study introduces FC3, a single, rG4-selective small molecule, as a minimal, mechanism-driven inducer and maintainer of totipotent stem cells (mG4TotiSCs and hG4TotiSCs). FC3 operates through a unified molecular mechanism and direct resolution of conserved rG4 structures in GATA2 mRNA, to elevate GATA2 protein synthesis and license the totipotency program. Given the evolutionary conservation of this G4 motif across mammals, G4 unwinding represents a conserved molecular strategy for robust in vitro induction of totipotent stem cells. Therefore, this method provides a robust platform applicable across stem cell research, disease modelling, assisted reproductive technologies, and fundamental studies of early mammalian development.

Over the past two decades, research into the biological roles of G4s has been overwhelmingly concentrated on pathological contexts, including oncogenesis, neurodegenerative disorders, and viral replication^45–48^, largely due to the prevalence of G4-stabilizing ligands as therapeutic candidates in these fields. Critically, nearly all existing G4-targeting tools are stabilizers^26^, which lock G4 structures into rigid conformations to perturb transcription or replication, a strategy inherently unsuited for probing dynamic, regulatory G4 functions in development or cell fate control. Although recent advances have yielded selective G4 unwinders^33,34^, including PhpC and FC3, their functional biology remains profoundly underexplored: no study to date has linked a defined G4 unwinder to a specific endogenous mRNA target, quantified its impact on translation in situ, or connected it to a fundamental developmental transition. Here, we bridge this critical gap by demonstrating that FC3 directly resolves a conserved rG4 motif in the CDS of GATA2 mRNA, thereby enhancing ribosome occupancy, boosting GATA2 protein synthesis, and licensing the totipotency program in both mouse and human stem cells. This work establishes the first causal, mechanism-resolved paradigm for G4 unwinding in mammalian cell identity regulation, and repositions G4 unwinders not as mere chemical tools, but as precision epigenetic modulators with broad potential across regenerative medicine, including reprogramming of tumor-initiating cells, modulation of immune cell plasticity, and intervention in age-associated translational decline.

While this study identifies a previously unappreciated role for rG4 unwinding in totipotency acquisition, there are still several important limitations. Although our data strongly support Gata2 knockout abolishes FC3-induced reprogramming, the possibility of cooperative or parallel mechanisms cannot be excluded. Notably, prior work has shown that exogenous overexpression of GATA2 alone is insufficient to activate the totipotency transcriptome in mESCs^39^, indicating that elevated GATA2 is necessary but insufficient. Consistent with this, genome-wide rG4 mapping (**Fig. 1b**) reveals that numerous maternal, ZGA and core totipotency-associated genes harbor rG4 motifs. Thus, FC3 may coordinately enhance the translation of multiple rG4-regulated factors, including but not limited to GATA2, to establish a permissive molecular environment for totipotency induction. It is noteworthy that neither mTotiSCs nor hTotiSCs activate the canonical totipotency-associated DUX pathway (**Extended Data Fig. 10**), thereby uncovering FC3’s distinct induction mechanism. As illustrated in **Fig. 6l**, we propose a model wherein FC3-mediated global rG4 remodelling acts synergistically with GATA2 upregulation to remodel chromatin accessibility, reset transcriptional networks, and license totipotency. Rigorous identification of FC3’s full target RNA repertoire and functional validation of individual candidates will be essential to dissect this multilayered regulatory architecture.

In summary, this study demonstrated that FC3-mediated rG4 unwinding provides a minimal, chemically defined, and mechanistically understood strategy for the robust derivation and long-term maintenance of functionally validated totipotent stem cells from both mouse and human pluripotent precursors. Critically, it reveals an unexpected and essential regulatory function for rG4s, not merely as transcriptional roadblocks, but as dynamic translational rheostats that control the expression of key regulators such as GATA2 and coordinate the broader totipotency network. These findings provide a conceptual and practical basis for re-examining rG4 biology in contexts requiring precise and rapid translational control, such as early embryonic development, gametogenesis, and somatic cell reprogramming. While our work delineates a core axis, the differential impact of FC3 on pluripotency-associated versus totipotency-associated transcripts remains incompletely resolved; future studies integrating time-resolved rG4 mapping, nascent RNA sequencing, and single-cell multi-omics will be essential to elucidate how rG4 dynamics orchestrate stage-specific gene expression programs during cell fate transitions.

## Acknowledgements

This work was financially supported by the Beijing Life Science Academy Initiative Scientific Research Program (2024101RPIA02 to J.L.), the National Natural Science Foundation of China (22504136 to X.T., T2588102 to J.L., 8227071346 to H.L. and 22474133 to Y.D.), the National Key Research and Development Program of China (2021YFA1200104 to J.L.), the New Cornerstone Science Foundation, the Natural Science Foundation of Hunan Province of China (2024JJ2028 to H.L.) and. We thank Prof. Zijiang Chen from Shandong University for kindly providing hESC and EGFP-hESC cell lines. We thank Prof. Xiaohua Shen from Basic Medical School, Tsinghua University, and Prof. Wei Xie from School of Life Sciences, Tsinghua University for valuable discussion of this work. We thank the Laboratory Animal Research Center, Tsinghua University, and Laboratory Animal Research Center, University of Science and Technology of China, for assisting with mouse embryo microinjection and transplantation. We thank Bin Yu from the Core Facility, Center of Biomedical Analysis, Tsinghua University, for technical support with flow cytometry analysis.

## Author contributions

Y.D., X.T., J.C.I.B., H.L. and J.L. designed the study; Y.D. and Q.Z. performed the cellular and mouse experiments; and Y.D., X.T., Q.Z., R.L., H.L. and J.L. interpreted the data. X.T. performed the library preparation and bioinformatics analysis of the RNA-seq, scRNA-seq, ATAC-seq, CUT&Tag and RiboLace data. Y.W. and K.Q. assisted with the bioinformatics analysis. W.H. and N.Z. performed the NMR analysis. D.H., X.C., L.D. and J.X. assisted with parts of the experiments. X.T., H.L. and J.L. conceived this project. X.T., J.C.I.B., H.L. and J.L. supervised the study. Y.D., X.T., Q.Z., H.L. and J.L. wrote the manuscript with input from all the authors. J.L. is the lead contact.

