## Extended Data Figures for "Unwinding of RNA G-quadruplexes induces mouse and human totipotency"

**Extended Data Figure**

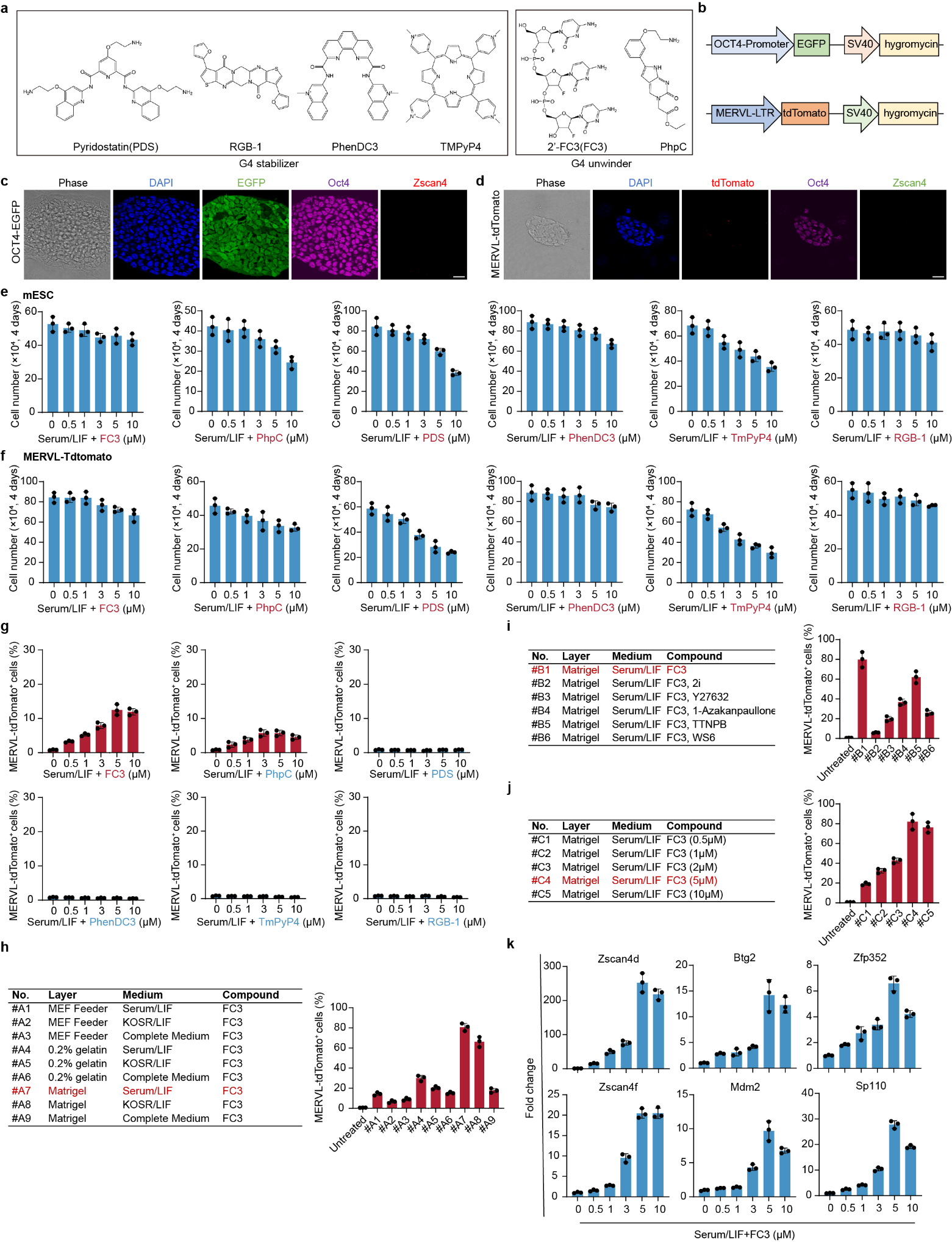

**Extended Data Figure 1 | Screening of G4 modulators that enable the induction of mouse totipotent stem cells and optimization of the induction conditions. a**, The chemical structures of 4 G4 stabilizers and 2 G4 unwinders. **b**, Schematic diagram of the OCT4-EGFP reporter gene and the MERVL-tdTomato reporter gene. **c**, Representative immunofluorescence images of OCT4 and ZSCAN4 in OCT4-EGFP mESCs. **d**, Representative immunofluorescence images of OCT4 and ZSCAN4 in MERVL-tdTomato mESCs. **e**, Proliferation of mESCs cultured on plates coated with Matrigel in serum/LIF medium supplemented with G4 stabilizers or unwinders at different concentrations for 4 days. The data are presented as the means ± s.d. of 3 independent experiments. **f**, Proliferation of MERVL-tdTomato mESCs cultured on plates coated with Matrigel in serum/LIF medium supplemented with G4 stabilizers or unwinders at different concentrations for 4 days. The data are presented as the means ± s.d. of 3 independent experiments. **g**, Bar graph showing the percentage of MERVL-tdTomato^+^ cells generated by treatment with G4 stabilizers or unwinders. The data are presented as the means ± s.d. of 3 independent experiments. **h**, Detailed list of cell culture conditions with different layers and media (left) and a bar graph showing the percentage of MERVL-tdTomato^+^ cells (right). The data are presented as the means ± s.d. of 3 independent experiments. **i**, Detailed list of cell culture conditions with different compounds (left) and bar graph showing the percentage of MERVL-tdTomato^+^ cells (right). The data are presented as the means ± s.d. of 3 independent experiments. **j**, Detailed list of the concentrations of FC3 (left) and bar graph showing the percentage of MERVL-tdTomato^+^ cells (right). The data are presented as the means ± s.d. of 3 independent experiments. **k**, RT-qPCR analysis of the relative expression of totipotent genes in mESCs treated with FC3 at different concentrations. The data were normalized to Gapdh. The data are presented as the means ± s.d. of 3 independent experiments.

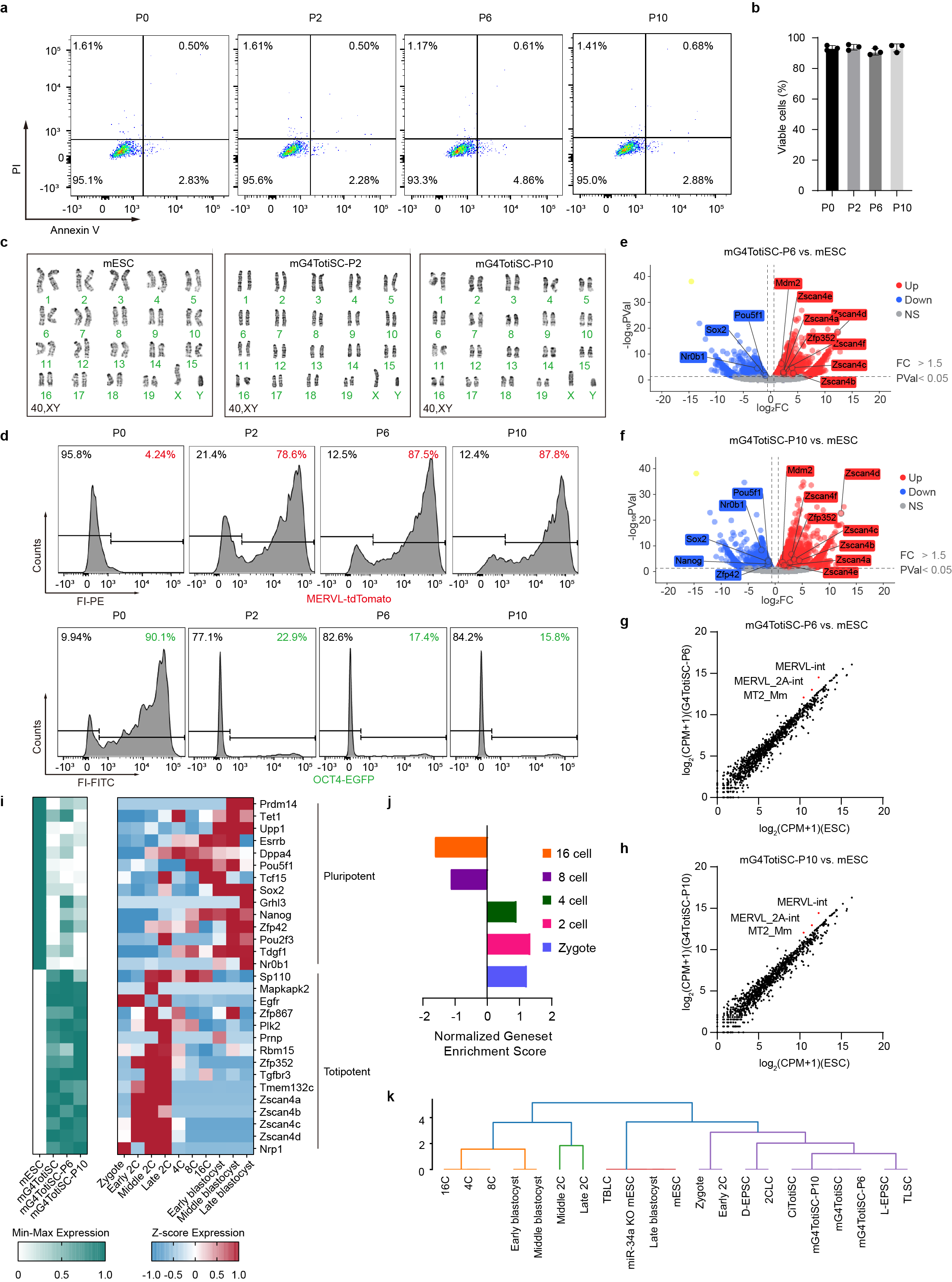

**Extended Data Figure 2 | FC3 enables the establishment and maintenance of G4TotiSCs in vitro, related to Figure 1. a**, Viability of mESCs (P0) and mG4TotiSCs (P2, P6, P10) was measured via Annexin V plus PI and analysed via flow cytometry. **b**, Bar graph showing the percentages of viable mESCs (P0) and mG4TotiSCs (P2, P6, P10). **c**, Karyotype analysis of mESCs (P0) and mG4TotiSCs (P2, P10). **d**, FACS analysis of mESCs (P0) or mG4TotiSCs (P2, P6, P10) from MERVL-tdTomato^+^ cells (top) and OCT4-EGFP^+^ cells (bottom). **e**, RNA-seq analysis of mESCs (P0) and G4TotiSCs (P6). The global transcriptomic changes are displayed in the volcano plot. Red and blue dots indicate upregulated (fold change ˃ 1.5) and downregulated (fold change ˂ 0.67) genes, respectively, with a p value < 0.05. **f**, RNA-seq analysis of mouse mESCs (P0) and G4TotiSCs (P10). The global transcriptomic changes are displayed in the volcano plot. Red and blue dots indicate upregulated (fold change ˃ 1.5) and downregulated (fold change ˂ 0.67) genes, respectively, with a p value < 0.05. **g**, Scatterplot displaying the transposon transcript comparison of mESCs (P0) and mG4TotiSCs (P6) via RNA-seq. **h**, Scatterplot displaying the transposon transcript comparison of mESCs (P0) and mG4TotiSCs (P10) via RNA-seq. **i**, Heatmap showing the relative expression of representative pluripotent and totipotent genes in mESCs, mG4TotiSCs and mouse embryos. **j**, GSEA of the bulk RNA-seq data from mG4TotiSCs according to the indicated embryonic stage-specific gene sets. **k**, Unsupervised hierarchical clustering analysis of mESCs, mG4TotiSCs, 2CLCs, CiTotiSCs, D-EPSCs, miR-24a KO mESCs, TBLCs, TLSCs and mouse embryos via RNA-seq.

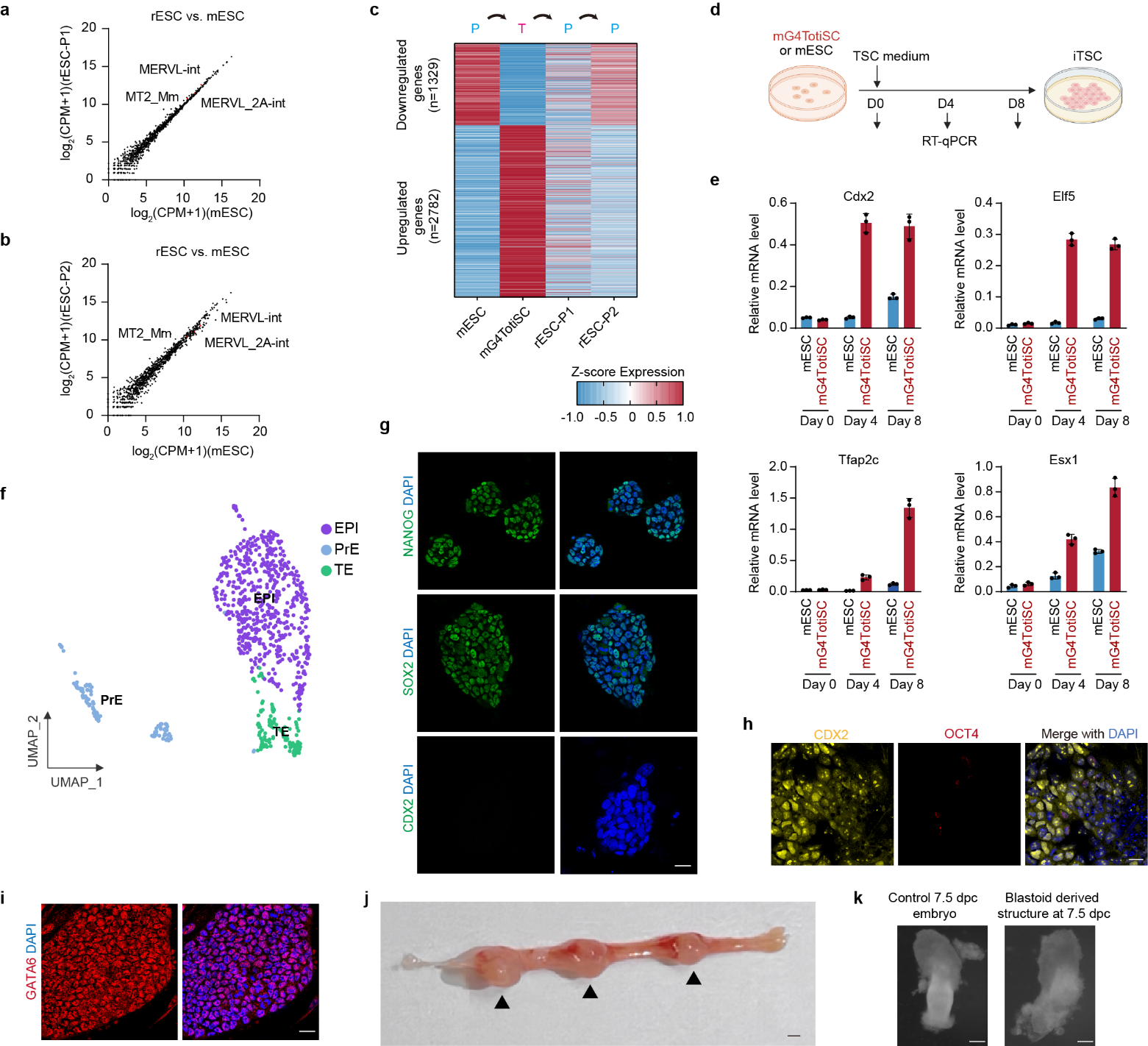

**Extended Data Figure 3 | Differentiation and self-organization ability of G4TotiSCs, related to Fig. 3**. **a**, Scatterplot displaying the transposon transcript comparison of rESCs (rP1) and mESCs via RNA-seq. **b**, Scatterplot displaying the transposon transcript comparison of rESCs (rP2) and mESCs via RNA-seq. **c**, Heatmap of up- and downregulated genes in mG4TotiSCs compared with mESCs and rESCs (rP1, rP2). **d**, Schematic diagram showing the differentiation and identification of TSCs from mESCs and mG4TotiSCs. **e**, RT‒qPCR analysis of the TSC markers Cdx2, Elf5, Tfap2c and Esx1 on days 0, 4 and 8 from mESCs and G4TotiSCs. The data are presented as the means ± s.d. of 3 independent experiments. **f**, Uniform manifold approximation and projection (UMAP) plot from the scRNA-seq data showing 3 clusters of mG4TotiSC-derived blastoids. **g**, Immunofluorescence imaging of NANOG, SOX2 and CDX2 expression in ESCs differentiating from mG4TotiSC-derived blastoids. Scale bar, 25 μm. **h**, Immunofluorescence imaging of OCT4 and CDX2 expression in TSCs differentiating from mG4TotiSC-derived blastoids. Scale bar, 25 μm. **i**, Immunofluorescence imaging of GATA6 expression in XEN cells differentiating from mG4TotiSC-derived blastoids. Scale bar, 25 μm. **j**, Image showing the formation of decidua in the mouse uterus 5 days after mG4TotiSC-derived blastoid transfer at 2.5 dpc. Black arrowheads indicate deciduae. Scale bar, 1 mm. **k**, Bright-field images of a control 7.5 dpc embryo or an in vivo mG4TotiSC-blastoid-derived structure recovered from decidua at 7.5 dpc (5 days after mG4TotiSC-derived blastoid transfer). Scale bar, 100 μm.

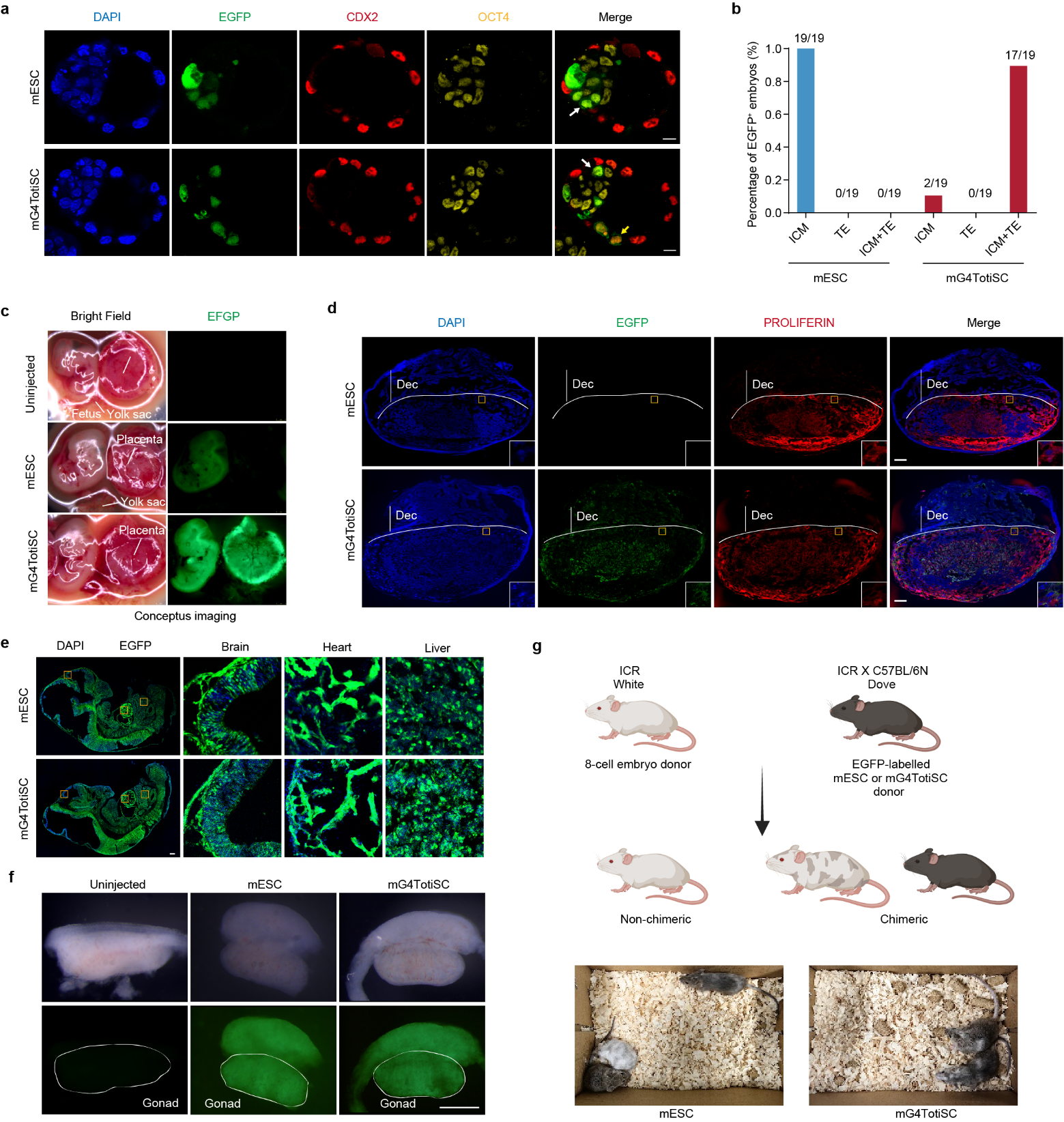

**Extended Data Figure 4 | Characterization of the chimerism potential of G4TotiSCs. a**, Representative immunofluorescence images showing the expression of CDX2 in chimeric blastocysts generated from 8-cell embryos injected with 3 EGFP^+^ mG4TotiSCs or mESCs in vitro. Scale bars, 20 μm. **b**, Fractions of chimeric blastocysts in which 3 injected EGFP^+^ G4TotiSCs or mESCs contributed to the ICM, TE or both. The fraction is indicated above each bar. **c**, Representative fluorescence images showing chimeras at E13.5. **d**, Example placenta sections from E13.5 chimeras generated from 8-cell embryos injected with multiple EGFP^+^ mG4TotiSCs or mESCs and stained with the trophoblast cell marker proliferin. Insets show enlarged images of single cells. Scale bars, 500 μm. **e**, Representative images of E13.5 chimeric embryo sections derived from 8-cell embryos injected with multiple EGFP^+^ mG4TotiSCs or mESCs. **f**, A representative image of E13.5 gonads from chimera contributed by mG4TotiSCs or mESCs. Scale bars, 500 μm. **g**, G4TotiSCs or mESC-derived chimeric mice.

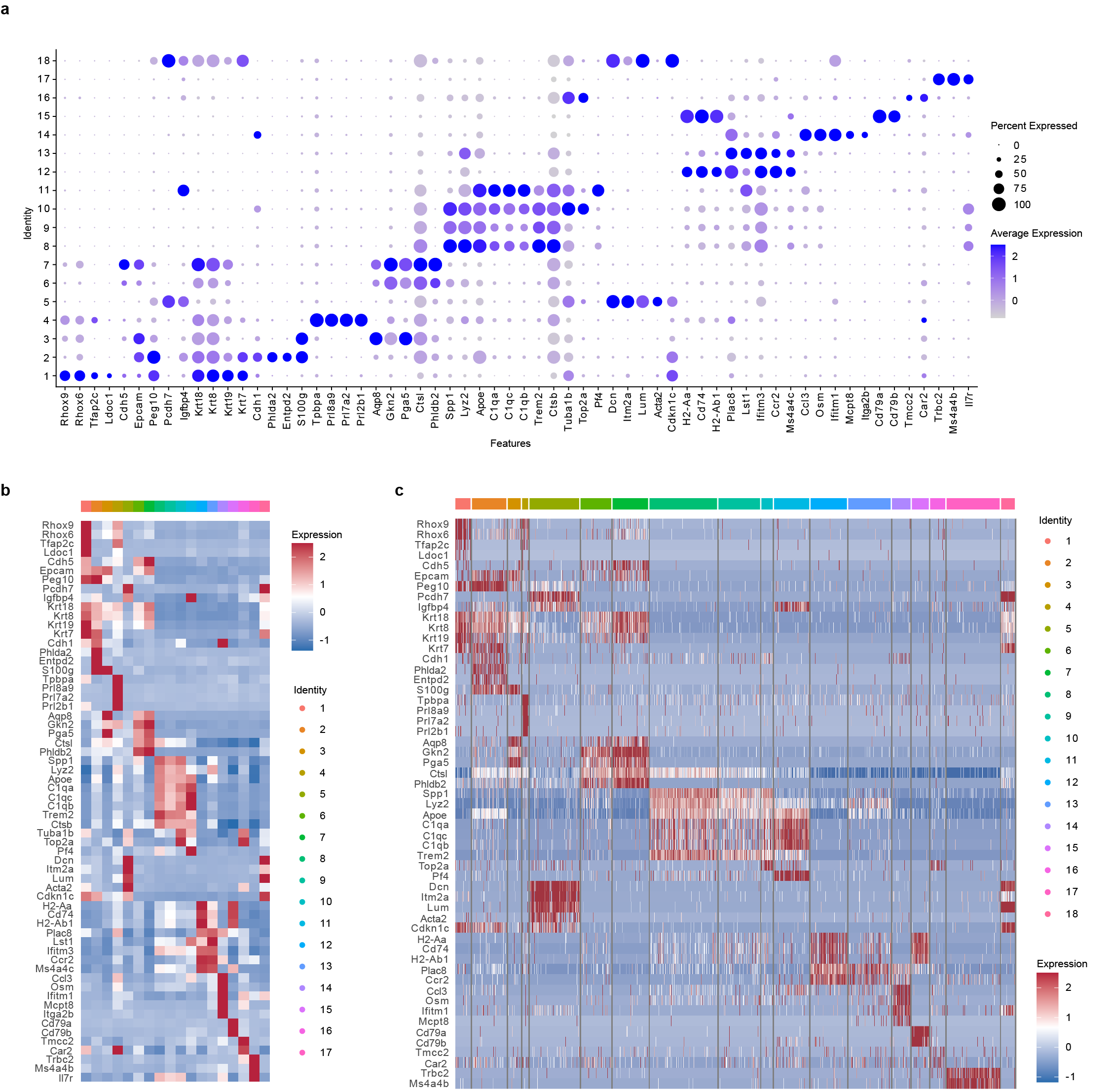

**Extended Data Figure 5 | ScRNA-seq analysis of the lineage contribution of the placenta and yolk sac derived from EGFP^+^ mG4TotiSCs via (a) bubble plots, (b) heatmaps and (c) heatmaps showing each cell.**

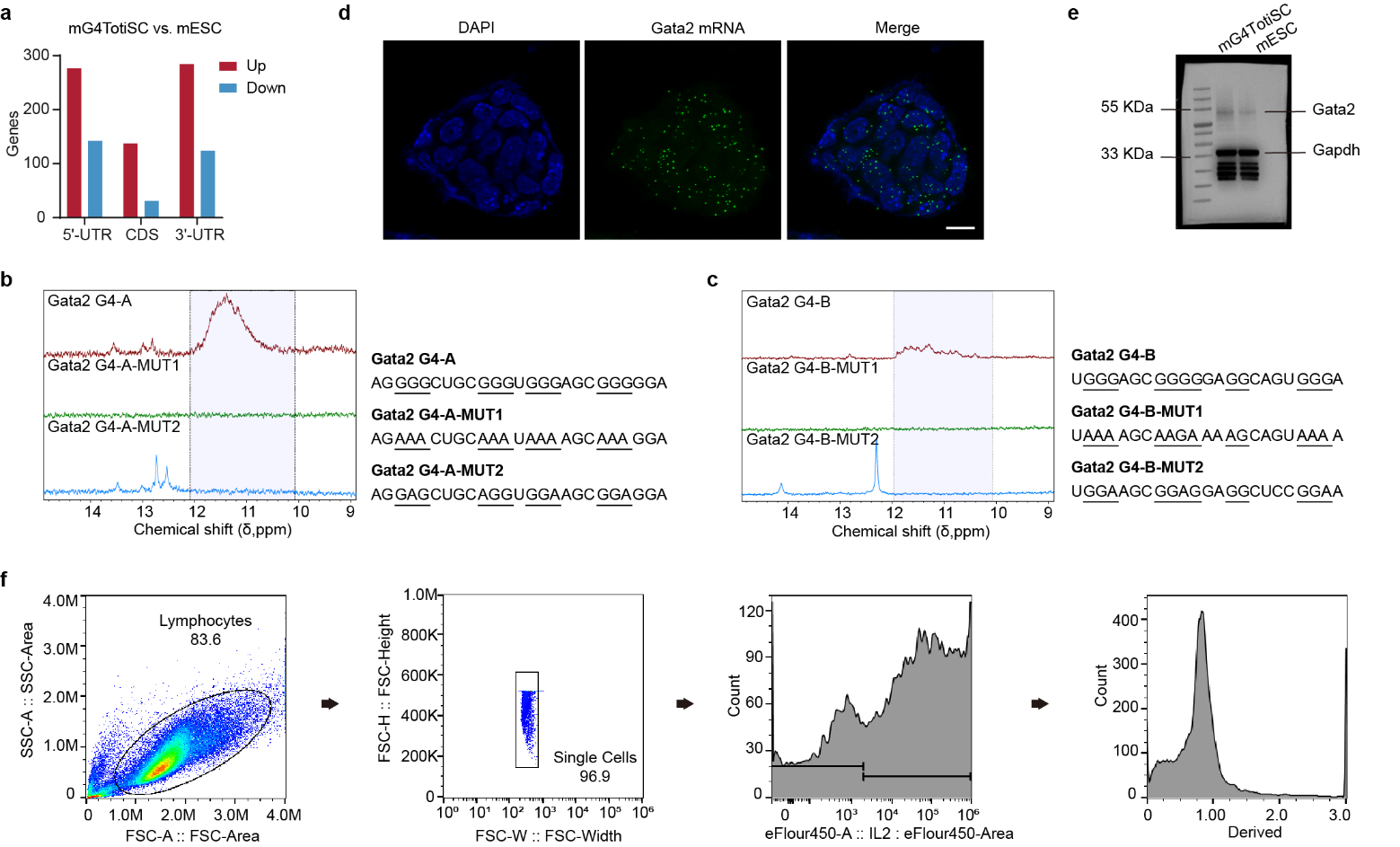

**Extended Data Figure 6 | Exploration of the mechanism by which FC3 induces totipotency transition. a**, Number of G4-containing mRNAs whose translation efficiency was upregulated or downregulated in mG4TotiSCs compared with that in mESCs. **b**, NMR analysis of the rG4 structure of the Gata2 G4-A oligo compared with the mutated Gata2 G4-A oligo. **c**, NMR analysis of the rG4 structure of the Gata2 G4-B oligo compared with the mutated Gata2 G4-B oligo. **d**, RCA imaging of intrinsic Gata2 mRNA expression in mESCs. Scale bar, 20 μm. **e**, Full-size image of the western blotting of GATA2 in mESCs and mG4TotiSCs. **f**, FACS gating strategy for Fig. 5f.

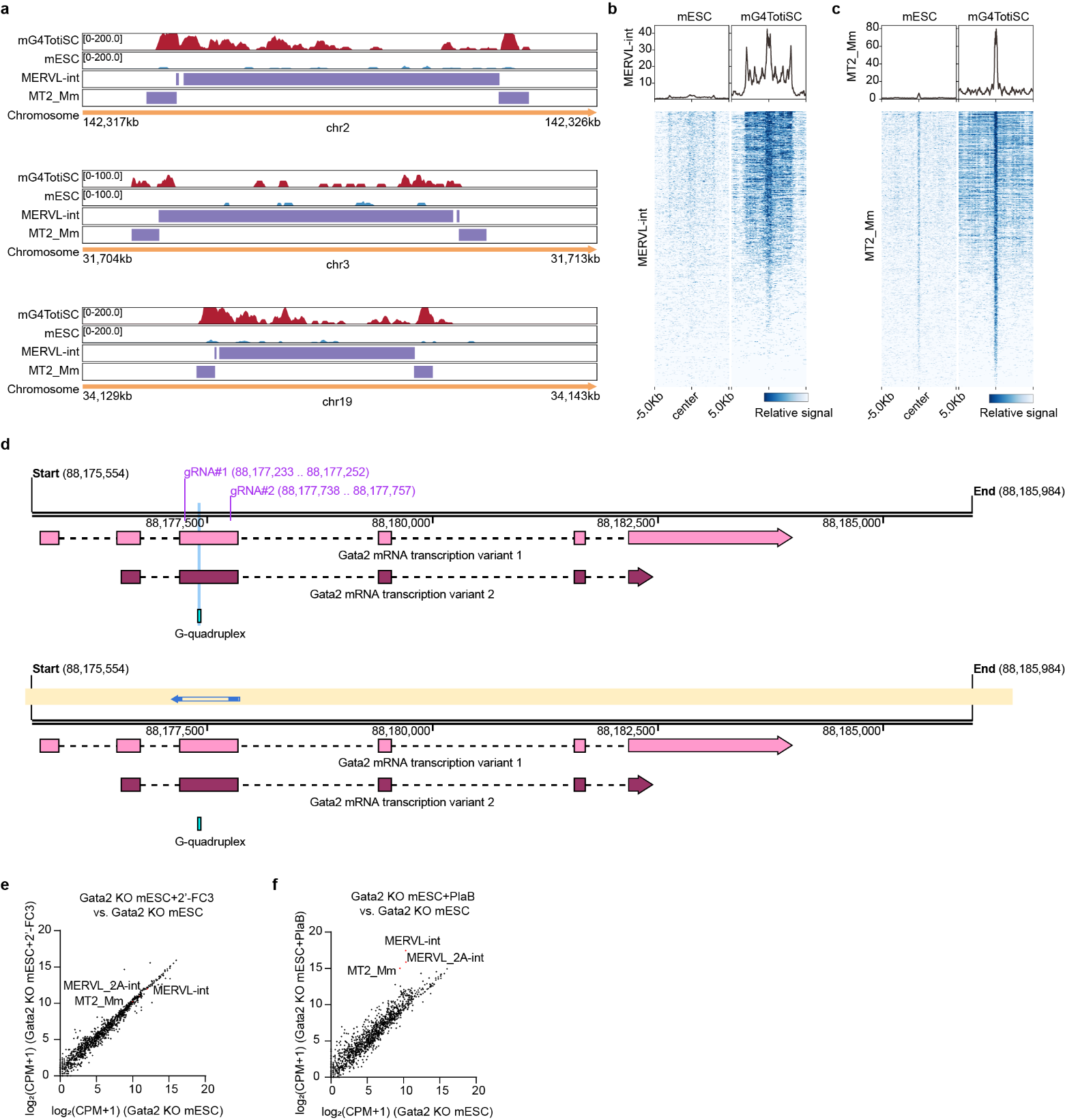

**Extended Data Figure 7 | FC3-induced totipotency transition is dependent on Gata2. a**, Gata2 cleavage under targets and tagmentation (CUT&Tag) peaks around MERVL-int and MT2-Mm transposons in mG4TotiSCs and mESCs. **b** and **c**, Heatmaps showing the Gata2 CUT&Tag peaks around the center of the MERVL-int and MT2-Mm transposons in mG4TotiSCs compared with mESCs. **d**, Schematic diagram of the experimental design and Sanger sequencing results for Gata2 KO mESCs. **e**, Scatterplots displaying the transposon transcript comparison of Gata2 KO mESC+2’-FC3 and Gata2 KO mESC via RNA-seq. Red dots, the key transposons MERVL-int, MERVL_2A-int, and MT2_Mm. **f**, Scatterplots displaying the transposon transcript comparison of Gata2 KO mESC+PlaB and Gata2 KO mESC via RNA-seq. Red dots, the key transposons MERVL-int, MERVL_2A-int, and MT2_Mm.

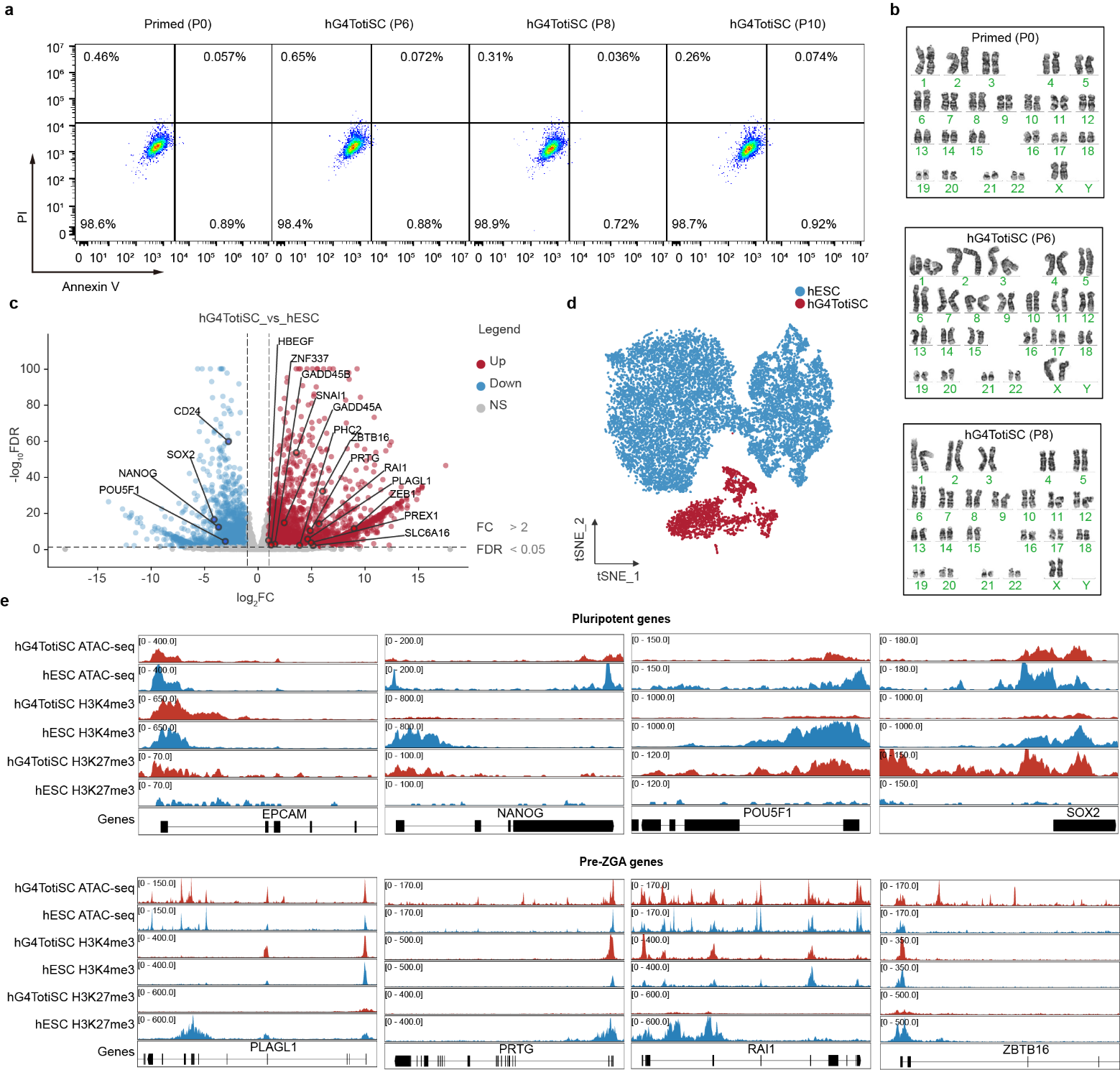

**Extended Data Figure 8 | Transcriptomic and epigenomic evaluation of the totipotent characteristics of hG4TotiSCs. a**, Viability of hESCs (P0) and hG4TotiSCs (P6, P8, P10) was measured via Annexin V plus PI and analysed via flow cytometry. **b**, Karyotype analysis of hESCs (P0) and hG4TotiSCs (P6, P8). **c**, RNA-seq analysis of mESCs and mG4TotiSCs. The global transcriptomic changes are displayed in the volcano plot. Red and blue dots indicate upregulated (fold change ˃ 2) and downregulated (fold change ˂ 2) genes, respectively, with a FRD ˂ 0.05. **d**, t-distributed stochastic neighbor embedding (t-SNE) plot from the scRNA-seq data showing 2 clusters of hESCs and hG4TotiSCs. **e**, ATAC-seq peaks, H3K4me3 and H3K27ac CUT&Tag peaks around representative pluripotent and totipotent genes in hG4TotiSCs and hESCs.

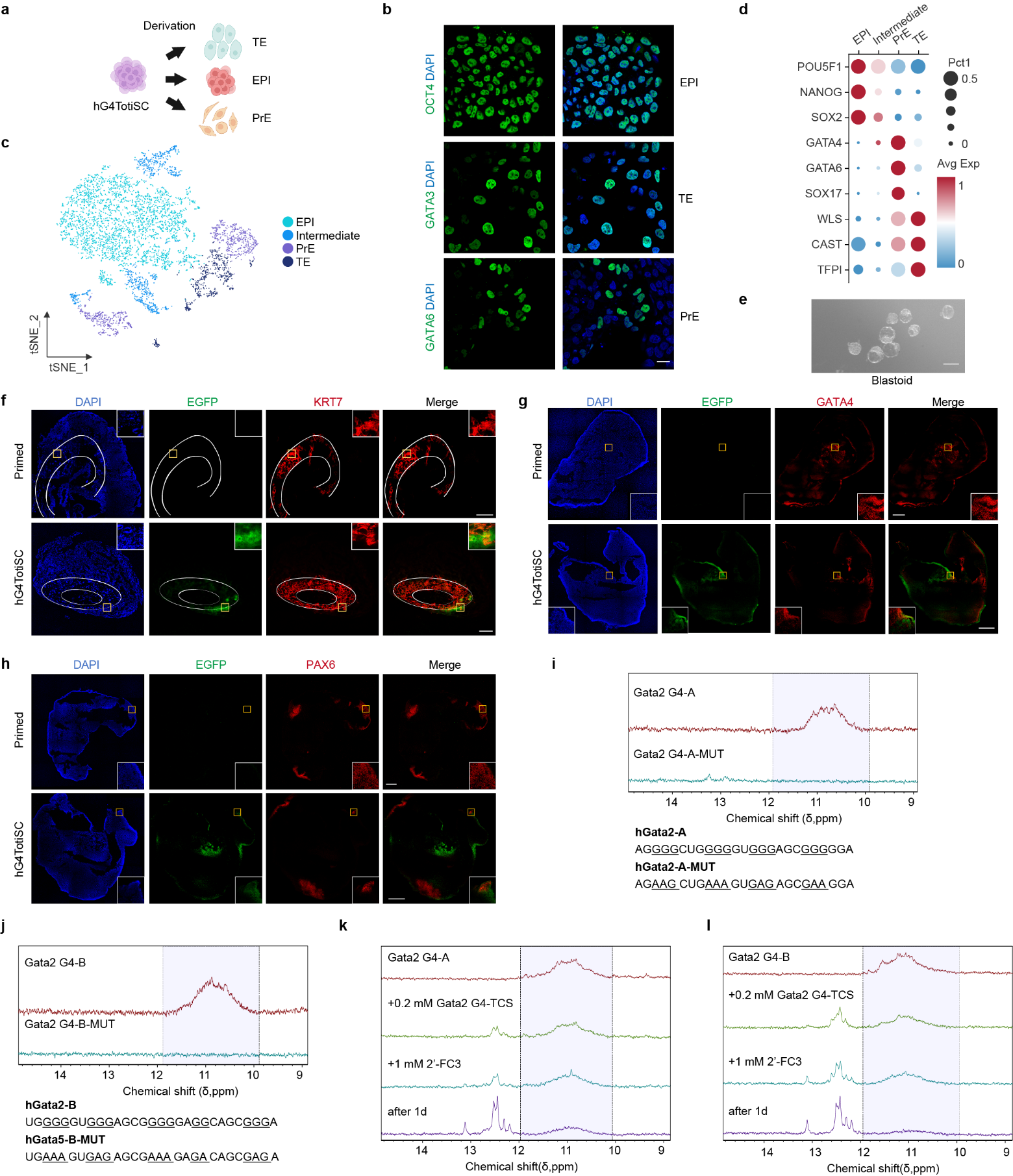

**Extended Data Figure 9 | hG4TotiSCs have bidirectional differentiation potential. a**, Schematic diagram showing the spontaneous lineage differentiation of hG4TotiSCs. **b**, Immunofluorescence imaging of OCT4, GATA3 or GATA6 expression in EPI, TE or PrE differentiating from hG4TotiSCs. Scale bar, 25 μm. **c**, t-SNE plot from the scRNA-seq data showing 4 clusters of EPI, intermediate, PrE and TE. **d**, Bubble plot from scRNA-seq showing the relative expression of representative EPI, PrE and TE genes in cells differentiating from hG4TotiSCs. **e**, Morphology of mG4TotiSC-derived blastoids. Scale bar, 100 μm. **f**, Representative immunofluorescence images showing the expression of KRT7 in E10.5 chimeras generated from 8-cell embryos injected with EGFP-labeled hG4TotiSCs or hESCs. Insets show enlarged images of single cells. Scale bars, 500 μm. **g**, Representative immunofluorescence images showing the expression of GATA4 in E10.5 chimeras generated from 8-cell embryos injected with EGFP-labeled hG4TotiSCs or hESCs. Scale bars, 500 μm. **h**, Representative immunofluorescence images showing the expression of PAX6 in E10.5 chimeras generated from 8-cell embryos injected with EGFP-labeled hG4TotiSCs or hESCs. Scale bars, 500 μm. **i**, NMR analysis of the rG4 structure of the hGata2 G4-A oligo compared with the mutated hGata2 G4-A oligo. **j**, NMR analysis of the rG4 structure of the hGata2 G4-B oligo compared with the mutated hGata2 G4-B oligo. **k**, NMR spectra of hGata2 G4-A oligos sequentially incubated with truncated complementary strands (TCSs) and FC3. **l**, NMR spectra of hGata2 G4-B oligos sequentially incubated with TCSs and FC3.

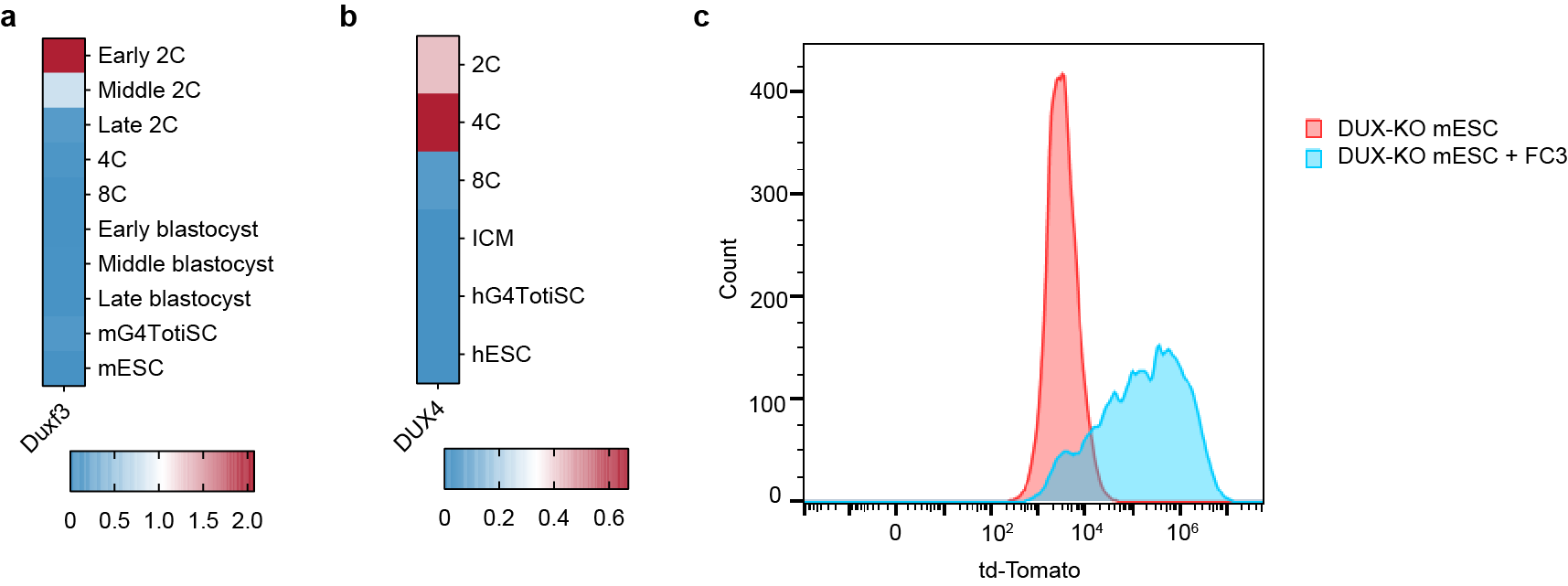

**Extended Data Figure 10 | mG4TotiSCs and hG4TotiSCs is independent on DUX pathway. a,** Heatmap showing expression levels of Duxf3 across mouse embryonic stages (early 2-cell to late blastocyst), mESCs, and mG4TotiSCs via RNA-seq. **b**, Heatmap showing expression levels of DUX4 across human embryonic stages (2-cell to ICM), hESCs, and hG4TotiSCs via RNA-seq. **c**, FC3 induces MERVL-driven TdTomato expression in DUX-KO MERVL-TdTomato mESCs. DUX-KO MERVL-TdTomato mESCs were treated with SLFM for 10 days and evaluated via FACS.
