## Supporting Information for "Unwinding of RNA G-quadruplexes induces mouse and human totipotency"

**Content**

**Table S1. The sources of all the chemicals, reagents, antibodies, cell lines, mice and commercial assay kits used**

| **Reagent or Resource** | **Source** | **Identifier** |
| --- | --- | --- |
| **Antibodies** | | |
| Oct3/4 Antibody (C-10) | Santa Cruz Biotechnology | Cat#sc-5279; RRID: AB_628051 |
| Anti-Zscan4 Antibody | Millipore | Cat#AB4340; RRID: AB_2827621 |
| MuERVL-Gag Polyclonal Antibody | Epigentek | Cat#A-2801; RRID: AB_3097813 |
| CDX2 Recombinant Rabbit Monoclonal Antibody | Thermo Fisher Scientific | Cat#MA5-35215; RRID: AB_2849119 |
| Anti-Oct4 antibody | Abcam | Cat#ab181557; RRID: AB_2687916 |
| Proliferin Antibody (E-10) | Santa Cruz Biotechnology | Cat#sc-271891; RRID: AB_10710396 |
| Rabbit anti-GAPDH polyclonal antibody | Yeasen | Cat#30202ES40; RRID: AB_2815025 |
| AffiniPure Goat Anti-Mouse IgG H&L/HRP | Bioss | Cat#bs-40296G-HRP; RRID: AB_3076686 |
| Goat Anti-Rabbit IgG H&L/HRP | Bioss | Cat#bs-0295G-HRP; RRID: AB_10923693 |
| Cy3-AffiniPure Goat Anti-mouse IgG(H+L) | Yeasen | Cat#33208ES60; RRID: AB_3105854 |
| YSFluor™ 647 Goat Anti-Rabbit lgG(H+L) | Yeasen | Cat#33113ES60; RRID: AB_3105856 |
| Donkey anti-Mouse lgG (H+L) Highly Cross-Adsorbed Secondary Antibody, Alexa Fluor^TM^ 555 | Thermo Fisher Scientific | Cat#A-31570; RRID: AB_2536180 |
| Donkey anti-Rabbit lgG (H+L) Highly Cross-Adsorbed Secondary Antibody, Alexa Fluor^TM^ 647 | Thermo Fisher Scientific | Cat#A-31573; RRID: AB_2536183 |
| Donkey anti-Mouse lgG (H+L) Highly Cross-Adsorbed Secondary Antibody, Alexa Fluor^TM^ 488 | Thermo Fisher Scientific | Cat#A-21202; RRID: AB_141607 |
| Donkey anti-Rat lgG (H+L) Cross-Adsorbed Secondary Antibody, DyLight^TM^ 755 | Thermo Fisher Scientific | Cat#SA5-10031; RRID: AB_2556611 |
| OCT3/4 Monoclonal Antibody (EM92), eBioscience | Thermo Fisher Scientific | Cat#14-5841-82; RRID: AB_914301 |
| ZSCAN4 Monoclonal Antibody (OTI1A6) | Thermo Fisher Scientific | Cat#MA5-26458; RRID: AB_2723809 |
| Goat Anti-Rabbit IgG Fc | Abcam | Cat#ab97196; RRID: AB_10680931 |
| GATA-2 (E9T6F) Rabbit mAb | Cell Signaling Technology | Cat#79802S; RRID: AB_3105938 |
| Pax6 (D3A9V) Rabbit Monoclonal Antibody | Cell Signaling Technology | Cat#60433T; RRID:AB_2797599 |
| Anti-GATA4 Antibody | Abcam | Cat#ab307823; RRID:AB_3105880 |
| KRT7 Rabbit mAb | Abclonal | Cat#A4357; RRID:AB_2863248 |
| Human GATA3 Antibody | R&D Systems | Cat#AF2605; RRID:AB_2108571 |
| ELF5 Polyclonal Antibody | Invitrogen | Cat#PA5-77158; RRID:AB_272088 |
| GATA-6 (D61E4) Rabbit Monoclonal Antibody | Cell Signaling Technology | Cat#5851S; RRID:AB_10705521 |
| Anti-TBR2 / EOMES Antibody | Abcam | Cat#AB183991; RRID:AB_2721040 |
| Nanog Monoclonal Antibody | Proteintech | Cat#67255-1-lg; RRID:AB_2882529 |
| Anti-SOX2 Antibody | Abcam | Cat#ab171380; RRID:AB_2732072 |
| Anti-Plzf Antibody | Abcam | Cat#ab104854; RRID:AB_10710087 |
| ZSCAN4 Polyclonal Antibody | Invitrogen | Cat#PA5-20901; RRID:AB_11152831 |
| **Chemicals, buffers and enzymes** | | |
| DMEM | Gibco | Cat#11995065 |
| MEM Non-Essential Amino Acids Solution (100*) | Gibco | Cat#11140035 |
| KnockOut^TM^ DMEM | Gibco | Cat#10829018 |
| RPMI Medium 1640 basic (1*) | Gibco | Cat#11875093 |
| Penicillin‒Streptomycin (100*) | Yeasen | Cat#60162ES76 |
| Gelatin, Type A | Merck | Cat#924504 |
| Fetal Bovine Serum | Gibco | Cat#30044333 |
| GlutaMAX^TM^ | Gibco | Cat#35050061 |
| Sodium pyruvate (100 mM) | Gibco | Cat#11360070 |
| 2-Mercaptoethanol | Gibco | Cat#21985023 |
| ESGRO® Leukemia Inhibitory Factor (LIF) | Millipore | Cat#ESG1107 |
| Corning® Matrigel® hESC-Qualified Matrix, LDEV-free | Corning | Cat#354277 |
| Heparin | Sigma‒Aldrich | Cat#H3149 |
| KnockOut^TM^ Serum Replacement | Gibco | Cat#10828028 |
| CD lipid concentrate | Gibco | Cat#11905031 |
| Sodium L-ascorbyl-2-phosphate | Selleck | Cat#S5115 |
| 1-Azakenpaullone | Selleck | Cat#S7193 |
| WS6 | Selleck | Cat#S7442 |
| TTNPB | Selleck | Cat#S4627 |
| Mitomycin C | Inalco | Cat#1758-9327 |
| Opti-MEM^TM^ | Gibco | Cat#31985070 |
| FGF4 | R&D systems | Cat#235-F4 |
| Pyridostatin (PDS) | MCE | Cat#HY-15176 |
| RGB-1 | Greact | Cat#GT403 |
| PhenDC3 | MCE | Cat#HY-15594 |
| TMPyP4 | MCE | Cat#HY-108477 |
| PhpC | Greact | Cat#GT401 |
| 2'-F C3 (FC3) | Greact | Cat#GT402 |
| PD0325901 | Solarbio | Cat#IP1310 |
| CHIR99021 | Sigma‒Aldrich | Cat#SML1046 |
| Blasticidin S | Yeasen | Cat#60218ES10 |
| Pladienolide B | Abmole | Cat#M7145 |
| HEPES Free Acid (1 M) Buffer Solution | Yeasen | Cat#60117ES60 |
| Hieff Trans® mRNA Transfection Reagent | Yeasen | Cat#40809ES03 |
| Lentivirus Concentration Solution | Yeasen | Cat#41101ES50 |
| Polybrene (hexadimethrine bromide) | Yeasen | Cat#40804ES76 |
| DMEM/F-12 | Yeasen | Cat#11320033 |
| Mineral oil | Sigma‒Aldrich | Cat#M8410 |
| EmbryoMax M2 medium | Millipore | Car#MR-015-D |
| EmbryoMax KSOM Embryo Culture | Millipore | Cat#MR-020P-5F |
| Lipofectamine™ 3000 transfection reagent | Invitrogen | Cat#L3000015 |
| Fast Giemsa Stain | Yeasen | Cat#40751ES01 |
| Colchicine | Yeasen | Cat#54659ES70 |
| 4% Paraformaldehyde Fix Solution (PFA) | Yeasen | Cat#60536ES60 |
| Triton^TM^ X-100 | Sigma‒Aldrich | Cat#T8787 |
| Tween-20 | Sango Biotech | Cat#A100777 |
| PBS (1*) | Yeasen | Cat#41403ES76 |
| Trypsin (0.25%), phenol red | Gibco | Cat#15050065 |
| Trypsin (2.5%) | Gibco | Cat#15090046 |
| Anti-Adherence Rinsing Solution | Stemcell | Cat#07010 |
| 5* Hieff Canace PCR Master Mix (for Multiplex PCR) | Yeasen | Cat#10137ES08 |
| Normal Donkey Serum | Yeasen | Cat#36116ES10 |
| Bovine Serum Albumin V | Solarbio | Cat#A8020 |
| Prolong® Gold Antifade Reagent with DAPI | Cell Signaling Technology | Cat#8961S |
| Auto Fluo Quencher | Applygen | Cat#C1212 |
| TRIzol™ reagent | Invitrogen | Cat#15596018CN |
| UltraPure^TM^ Distilled Water | Invitrogen | Cat#10977015 |
| Ethanol | Macklin | Cat#E821483 |
| EDTA (0.5 M), pH 8.0, RNase free | Invitrogen | Cat#AM9260G |
| KCl (1 M, RNase free) | Feimobio | Cat#FB9997-500 |
| Dithiothreitol (DTT) | Inalco SpA | Cat#1758-9030 |
| NP-40 | Solarbio | Cat#N8030 |
| O.C.T. Compound | Yeasen | Cat#36309ES61 |
| HBSS (with Mg^2+^, Ca^2+^) | Yeasen | Cat#60147ES76 |
| StemPro^TM^ Accutase^TM^ | Gibco | Cat#A1110501 |
| Collagenase IV | Yeasen | Cat#40510ES60 |
| DNase I (Rnase-Free) | Tiangen | Cat#RT411 |
| Alamar Blue | Yeasen | Cat#40202ES76 |
| HEPES-KOH(1 M, pH 7.9) | Twbio | Cat#X0460 |
| Spermidine | Sigma‒Aldrich | Cat#85558-1G |
| MgCl_2_ (1 M, RNase free) | Leagene | Cat#NR0220 |
| Glycerol | Biorigin | Cat#BN20148-500 ml |
| Tn5 transposome | Greact | Cat#GTX01 |
| SDS (10% solution, Sodium Dodecyl Sulfate) | Invitrogen | Cat#AM9822 |
| Tagment DNA Extraction Beads | Novoprotein | Cat#N245-01B |
| Hieff NGS® DNA Selection Beads | Yeasen | Cat#12601ES56 |
| Sucrose | Yeasen | Cat#60350ES80 |
| Murine RNase Inhibitor | Vazyme | Cat#R301-03 |
| UltraPure™ 1 M Tris-HCl | Invitrogen | Cat#15567027 |
| 50*TAE Buffer | Yeasen | Cat#60116ES76 |
| YeaRed Nucleic Acid Gel Stain (10,000* in DMSO) | Yeasen | Cat#10203ES76 |
| 3:1Agarose | Yeasen | Cat#10220ES08 |
| Dynabeads^TM^ M-280 Streptavidin | Thermo Fisher Scientific | Cat#11206D |
| PMSF | Yeasen | Cat#58076ES01 |
| 2×KunGre ™ qPCR SYBR Green Mix | Greact | Cat#GT133-02 |
| Lysis Buffer for WB/IP Assays (No inhibitors) | Yeasen | Cat#20119ES60 |
| GoldBand Plus 3-color Range Protein Marker(8-180 kDa) | Yeasen | Cat#20350ES90 |
| Precast Protein Plus Gel, 4-20%, 15 wells | Yeasen | Cat#36256ES10 |
| Precast Running Buffer, 2 L (Powder) | Yeasen | Cat#36257ES05 |
| 5*SDS‒PAGE Protein Loading Buffer | Yeasen | Cat#20315ES20 |
| 10* TBST Buffer | Yeasen | Cat#60145ES80 |
| Western First Antibody Diluent Buffer | Beyotime | Cat#P0023A |
| Western Second Antibody Diluent Buffer | Beyotime | Cat#P0023D |
| Skim Milk | Yeasen | Cat#36120ES76 |
| Amersham^TM^ Protran^TM^ 0.45 NC | Cytiva | Cat#10600002 |
| dNTP Set Solution | Yeasen | Cat#10122ES74 |
| mTeSR^TM^1 medium | Stemcell | Cat#85850 |
| Dispase in DMEM/F-12 | Stemcell | Cat#07923 |
| Accumax | Thermo Fisher Scientific | Cat#00-4666-56 |
| Minocycline | MCE | Cat#HY-17412A |
| Versene | Gibco | Cat#15040066 |
| DNAse I Solution | Stemcell | Car#07900 |
| N-2 Supplement | Gibco | Cat#A1370701 |
| B-27 Supplement | Gibco | Cat#17504044 |
| Neurobasal^TM^ Medium | Gibco | Cat#21103049 |
| rhFGF-4 Protein | Yeasen | Cat#91303ES08 |
| Heparin | Yeasen | Cat#738416ES60 |
| TrypLE^TM^ Express | Gibco | Cat#12605010 |
| BMP-4 Protein | Yeasen | Cat#92070ES05 |
| A 83-01 | Yeasen | Cat#53002ES03 |
| M16 Medium | Sigma | Cat#MR-010 |
| G-2^TM^ plus | Vitrolife | Cat#10132 |
| **Critical commercial assays** | | |
| MolPure Cell/Tissue DNA Kit | Yeasen | Cat#18700ES70 |
| RNA Easy Fast Tissue/Cell Kit | Tiangen | Cat#DP451 |
| EndoFree Mini Plasmid Kit | Tiangen | Cat#DP118 |
| KunGre ™ 1st Strand cDNA Synthesis Kit for qPCR(gDNA digester plus) | Greact | Cat#GT141-02 |
| TIANgel Purification Kit | Tiangen | Cat#DP219 |
| Annexin V-PE/7-AAD cell apoptosis detection kit | Yeasen | Cat#40310ES20 |
| BCA Protein Quantification Kit | Yeasen | Cat#20201ES86 |
| Enhanced ECL Chemiluminescent Substrate Kit | Yeasen | Cat#36222ES60 |
| NEBNext® UltraTM RNA Library Prep Kit for Illumina | NEB | Cat#E7530L |
| Chromium Next GEM Single Cell 3ʹ Kit v3.1 | 10X Genomics | Cat#PN-1000268 |
| Chromium Next GEM Chip G Single Cell Kit | 10X Genomics | Cat#PN-1000127 |
| Zymo-Seq RRBS Library Kit | Zymo | Cat#D5460 |
| MolLink® High Quality CUT&Tag kit | Greact | Cat#GTX02 |
| TruePrep Index Kit V2 for Illumina | Vazyme | Cat#TD202 |
| 1*dsDNA HS Assay Kit | Yeasen | Cat#12642ES60 |
| AggreWell^TM^ 400 | Stemcell | Cat#34450 |
| **Cell lines** | | |
| Mouse cell line: mESC | This study | N/A |
| Mouse cell line: EGFP-labelled mESC | This study | N/A |
| Mouse cell line: OCT4-EGFP mESC | This study | N/A |
| Mouse cell line: MERVL-tdTomato mESC | This study | N/A |
| Mouse cell line: Gata2 KO mESC | This study | N/A |
| Mouse cell line: MERVL-TdTomato Dux-KO mESC | Gift from Prof. Wei Xie | N/A |
| Human cell line: HEK293FT | National Infrastructure of Cell Line Resource (NICLR, China) | N/A |
| Human cell line: hESC | Gift from Prof. Zijiang Chen | N/A |
| Human cell line: EGFP-hESC | Gift from Prof. Zijiang Chen | N/A |
| **Mouse strains** | | |
| ICR | Beijing Vital River Laboratory Animal Technology | Stock No: 201 |
| ICR-Tg(CAG-EGFP) 5Vst/Vst | Beijing Vitalstar Biotechnology | N/A |
| C57BL/6NCrl OCT4-EGFP | Beijing Vitalstar Biotechnology | N/A |

**Table S2. Plasmid sequences used in this work**

| **Plasmid** | **Sequence** | **Note** |
| --- | --- | --- |
| pLenti-MERVL-tdTomato | ACGCGTGTAGTCTTATGCAATACTCTTGTAGTCTTGCAACATGGTAACGATGAGTTAGCAACATGCCTTACAAGGAGAGAAAAAGCACCGTGCATGCCGATTGGTGGAAGTAAGGTGGTACGATCGTGCCTTATTAGGAAGGCAACAGACGGGTCTGACATGGATTGGACGAACCACTGAATTGCCGCATTGCAGAGATATTGTATTTAAGTGCCTAGCTCGATACAATAAACGGGTCTCTCTGGTTAGACCAGATCTGAGCCTGGGAGCTCTCTGGCTAACTAGGGAACCCACTGCTTAAGCCTCAATAAAGCTTGCCTTGAGTGCTTCAAGTAGTGTGTGCCCGTCTGTTGTGTGACTCTGGTAACTAGAGATCCCTCAGACCCTTTTAGTCAGTGTGGAAAATCTCTAGCAGTGGCGCCCGAACAGGGACCTGAAAGCGAAAGGGAAACCAGAGCTCTCTCGACGCAGGACTCGGCTTGCTGAAGCGCGCACGGCAAGAGGCGAGGGGCGGCGACTGGTGAGTACGCCAAAAATTTTGACTAGCGGAGGCTAGAAGGAGAGAGATGGGTGCGAGAGCGTCAGTATTAAGCGGGGGAGAATTAGATCGCGATGGGAAAAAATTCGGTTAAGGCCAGGGGGAAAGAAAAAATATAAATTAAAACATATAGTATGGGCAAGCAGGGAGCTAGAACGATTCGCAGTTAATCCTGGCCTGTTAGAAACATCAGAAGGCTGTAGACAAATACTGGGACAGCTACAACCATCCCTTCAGACAGGATCAGAAGAACTTAGATCATTATATAATACAGTAGCAACCCTCTATTGTGTGCATCAAAGGATAGAGATAAAAGACACCAAGGAAGCTTTAGACAAGATAGAGGAAGAGCAAAACAAAAGTAAGACCACCGCACAGCAAGCGGCCACTGATCTTCAGACCTGGAGGAGGAGATATGAGGGACAATTGGAGAAGTGAATTATATAAATATAAAGTAGTAAAAATTGAACCATTAGGAGTAGCACCCACCAAGGCAAAGAGAAGAGTGGTGCAGAGAGAAAAAAGAGCAGTGGGAATAGGAGCTTTGTTCCTTGGGTTCTTGGGAGCAGCAGGAAGCACTATGGGCGCAGCCTCAATGACGCTGACGGTACAGGCCAGACAATTATTGTCTGGTATAGTGCAGCAGCAGAACAATTTGCTGAGGGCTATTGAGGCGCAACAGCATCTGTTGCAACTCACAGTCTGGGGCATCAAGCAGCTCCAGGCAAGAATCCTGGCTGTGGAAAGATACCTAAAGGATCAACAGCTCCTGGGGATTTGGGGTTGCTCTGGAAAACTCATTTGCACCACTGCTGTGCCTTGGAATGCTAGTTGGAGTAATAAATCTCTGGAACAGATTGGAATCACACGACCTGGATGGAGTGGGACAGAGAAATTAACAATTACACAAGCTTAATACACTCCTTAATTGAAGAATCGCAAAACCAGCAAGAAAAGAATGAACAAGAATTATTGGAATTAGATAAATGGGCAAGTTTGTGGAATTGGTTTAACATAACAAATTGGCTGTGGTATATAAAATTATTCATAATGATAGTAGGAGGCTTGGTAGGTTTAAGAATAGTTTTTGCTGTACTTTCTATAGTGAATAGAGTTAGGCAGGGATATTCACCATTATCGTTTCAGACCCACCTCCCAACCCCGAGGGGACCCGACAGGCCCGAAGGAATAGAAGAAGAAGGTGGAGAGAGAGACAGAGACAGATCCATTCGATTAGTGAACGGATCTCGACGGTATCGGTTAACTTTTAAAAGAAAAGGGGGGATTGGGGGGTACAGTGCAGGGGAAAGAATAGTAGACATAATAGCAACAGACATACAAACTAAAGAATTACAAAAACAAATTACAAAATTCAAAATTTTATCGATgctagctgtagtggttattcctggttgtcaacttgacaatatttggaatgaactacagtccggaattggaaggctcaccagtgacccttatctggaggcttggagatccttatctggatcttggtttgaagatcttgagccatagtggctatggattccagaagattgaatctccgagtttaaggaacacacctttaatctgggctacgcctttcatctgggattaaaggtgtggtggaacacacctttaatctgggctacaccttctgctggagacaatataaggacattggaagaagggagtctagctcttgctcctgctccttcgcctgcttgctgcgtgagactgagtaactgctagatccttggacttccattcacagctgtgactgaacgattgttgggaattgagctgccgactgtaagtcatcaataaattcctttattatctagagactatccataagttctgtgactctagagaaccctgactaatacagaagttggtaccaggagtggttctagagtaacagaagtacaaggatgaatcttttaaaattctggaattggcttgttgatccaccagcactttcaactattgaaacctctccagattctctccctcctgggagctcagagaattttgaagacccatggttgaaactatattccgaacttaaagaagctaatgcccttgattttcttaatgaattaggtgattcagtgcacaaagcttggtaccgagctcggatccactagtaacggccgccagtgtgctggaattcgcccttgtctgtctagaatggtgagcaagggcgaggacgtcatcaaagagttcatgcgcttcaaggtgcgcatggagggctccatgaacggccacgagttcgagatcgagggcgagggcgagggccgcccctacgagggcacccagaccgccaagctgaaggtgaccaagggcggccccctgcccttcgcctgggacatcctgtccccccagttcatgtacggctccaaggcgtacgtgaagcaccccgccgacatccccgattacaagaagctgtccttccccgagggcttcaagtgggagcgcgtgatgaacttcgaggacggcggtctggtgaccgtgacccaggactcctccctgcaggacggcacgctgatctacaaggtgaagatgcgcggcaccaacttcccccccgacggccccgtaatgcagaagaagaccatgggctgggaggcctccaccgagcgcctgtacccccgcgacggcgtgctgaagggcgagatccaccaggccctgaagctgaaggacggcggccactacctggtggagttcaagaccatctacatggccaagaagcccgtgcaactgcccggctactactacgtggacaccaagctggacatcacctcccacaacgaggactacaccatcgtggaacagtacgagcgctccgagggccgccaccacctgttcctggggcatggcaccggcagcaccggcagcggcagctccggcaccgcctcctccgaggacaacaacatggccgtcatcaaagagttcatgcgcttcaaggtgcgcatggagggctccatgaacggccacgagttcgagatcgagggcgagggcgagggccgcccctacgagggcacccagaccgccaagctgaaggtgaccaagggcggccccctgcccttcgcctgggacatcctgtccccccagttcatgtacggctccaaggcgtacgtgaagcaccccgccgacatccccgattacaagaagctgtccttccccgagggcttcaagtgggagcgcgtgatgaacttcgaggacggcggtctggtgaccgtgacccaggactcctccctgcaggacggcacgctgatctacaaggtgaagatgcgcggcaccaacttcccccccgacggccccgtaatgcagaagaagaccatgggctgggaggcctccaccgagcgcctgtacccccgcgacggcgtgctgaagggcgagatccaccaggccctgaagctgaaggacggcggccactacctggtggagttcaagaccatctacatggccaagaagcccgtgcaactgcccggctactactacgtggacaccaagctggacatcacctcccacaacgaggactacaccatcgtggaacagtacgagcgctccgagggccgccaccacctgttcctgtacggcatggacgagctgtacaagtaaGCGGCCGCtaattctaccgggtaggggaggcgcttttcccaaggcagtctggagcatgcgctttagcagccccgctgggcacttggcgctacacaagtggcctctggcctcgcacacattccacatccaccggtaggcgccaaccggctccgttctttggtggccccttcgcgccaccttctactcctcccctagtcaggaagttcccccccgccccgcagctcgcgtcgtgcaggacgtgacaaatggaagtagcacgtctcactagtctcgtgcagatggacagcaccgctgagcaatggaagcgggtaggcctttggggcagcggccaatagcagctttgctccttcgctttctgggctcagaggctgggaaggggtgggtccgggggcgggctcaggggcgggctcaggggcggggcgggcgcccgaaggtcctccggaggcccggcattctgcacgcttcaaaagcgcacgtctgccgcgctgttctcctcttcctcatctccgggcctttcgacctgcagcccaagcttaccATGGCCAAGCCTTTGTCTCAAGAAGAATCCACCCTCATTGAAAGAGCAACGGCTACAATCAACAGCATCCCCATCTCTGAAGACTACAGCGTCGCCAGCGCAGCTCTCTCTAGCGACGGCCGCATCTTCACTGGTGTCAATGTATATCATTTTACTGGGGGACCTTGTGCAGAACTCGTGGTGCTGGGCACTGCTGCTGCTGCGGCAGCTGGCAACCTGACTTGTATCGTCGCGATCGGAAATGAGAACAGGGGCATCTTGAGCCCCTGCGGACGGTGCCGACAGGTGCTTCTCGATCTGCATCCTGGGATCAAAGCCATAGTGAAGGACAGTGATGGACAGCCGACGGCAGTTGGGATTCGTGAATTGCTGCCCTCTGGTTATGTGTGGGAGGGCTAAGTCGACAATCAACCTCTGGATTACAAAATTTGTGAAAGATTGACTGGTATTCTTAACTATGTTGCTCCTTTTACGCTATGTGGATACGCTGCTTTAATGCCTTTGTATCATGCTATTGCTTCCCGTATGGCTTTCATTTTCTCCTCCTTGTATAAATCCTGGTTGCTGTCTCTTTATGAGGAGTTGTGGCCCGTTGTCAGGCAACGTGGCGTGGTGTGCACTGTGTTTGCTGACGCAACCCCCACTGGTTGGGGCATTGCCACCACCTGTCAGCTCCTTTCCGGGACTTTCGCTTTCCCCCTCCCTATTGCCACGGCGGAACTCATCGCCGCCTGCCTTGCCCGCTGCTGGACAGGGGCTCGGCTGTTGGGCACTGACAATTCCGTGGTGTTGTCGGGGAAATCATCGTCCTTTCCTTGGCTGCTCGCCTGTGTTGCCACCTGGATTCTGCGCGGGACGTCCTTCTGCTACGTCCCTTCGGCCCTCAATCCAGCGGACCTTCCTTCCCGCGGCCTGCTGCCGGCTCTGCGGCCTCTTCCGCGTCTTCGCCTTCGCCCTCAGACGAGTCGGATCTCCCTTTGGGCCGCCTCCCCGCCTGGTACCTTTAAGACCAATGACTTACAAGGCAGCTGTAGATCTTAGCCACTTTTTAAAAGAAAAGGGGGGACTGGAAGGGCTAATTCACTCCCAACGAAAATAAGATCTGCTTTTTGCTTGTACTGGGTCTCTCTGGTTAGACCAGATCTGAGCCTGGGAGCTCTCTGGCTAACTAGGGAACCCACTGCTTAAGCCTCAATAAAGCTTGCCTTGAGTGCTTCAAGTAGTGTGTGCCCGTCTGTTGTGTGACTCTGGTAACTAGAGATCCCTCAGACCCTTTTAGTCAGTGTGGAAAATCTCTAGCAGTAGTAGTTCATGTCATCTTATTATTCAGTATTTATAACTTGCAAAGAAATGAATATCAGAGAGTGAGAGGAACTTGTTTATTGCAGCTTATAATGGTTACAAATAAAGCAATAGCATCACAAATTTCACAAATAAAGCATTTTTTTCACTGCATTCTAGTTGTGGTTTGTCCAAACTCATCAATGTATCTTATCATGTCTGGCTCTAGCTATCCCGCCCCTAACTCCGCCCAGTTCCGCCCATTCTCCGCCCCATGGCTGACTAATTTTTTTTATTTATGCAGAGGCCGAGGCCGCCTCGGCCTCTGAGCTATTCCAGAAGTAGTGAGGAGGCTTTTTTGGAGGCCTAGACTTTTGCAGAGACGGCCCAAATTCGTAATCATGGTCATAGCTGTTTCCTGTGTGAAATTGTTATCCGCTCACAATTCCACACAACATACGAGCCGGAAGCATAAAGTGTAAAGCCTGGGGTGCCTAATGAGTGAGCTAACTCACATTAATTGCGTTGCGCTCACTGCCCGCTTTCCAGTCGGGAAACCTGTCGTGCCAGCTGCATTAATGAATCGGCCAACGCGCGGGGAGAGGCGGTTTGCGTATTGGGCGCTCTTCCGCTTCCTCGCTCACTGACTCGCTGCGCTCGGTCGTTCGGCTGCGGCGAGCGGTATCAGCTCACTCAAAGGCGGTAATACGGTTATCCACAGAATCAGGGGATAACGCAGGAAAGAACATGTGAGCAAAAGGCCAGCAAAAGGCCAGGAACCGTAAAAAGGCCGCGTTGCTGGCGTTTTTCCATAGGCTCCGCCCCCCTGACGAGCATCACAAAAATCGACGCTCAAGTCAGAGGTGGCGAAACCCGACAGGACTATAAAGATACCAGGCGTTTCCCCCTGGAAGCTCCCTCGTGCGCTCTCCTGTTCCGACCCTGCCGCTTACCGGATACCTGTCCGCCTTTCTCCCTTCGGGAAGCGTGGCGCTTTCTCATAGCTCACGCTGTAGGTATCTCAGTTCGGTGTAGGTCGTTCGCTCCAAGCTGGGCTGTGTGCACGAACCCCCCGTTCAGCCCGACCGCTGCGCCTTATCCGGTAACTATCGTCTTGAGTCCAACCCGGTAAGACACGACTTATCGCCACTGGCAGCAGCCACTGGTAACAGGATTAGCAGAGCGAGGTATGTAGGCGGTGCTACAGAGTTCTTGAAGTGGTGGCCTAACTACGGCTACACTAGAAGGACAGTATTTGGTATCTGCGCTCTGCTGAAGCCAGTTACCTTCGGAAAAAGAGTTGGTAGCTCTTGATCCGGCAAACAAACCACCGCTGGTAGCGGTGGTTTTTTTGTTTGCAAGCAGCAGATTACGCGCAGAAAAAAAGGATCTCAAGAAGATCCTTTGATCTTTTCTACGGGGTCTGACGCTCAGTGGAACGAAAACTCACGTTAAGGGATTTTGGTCATGAGATTATCAAAAAGGATCTTCACCTAGATCCTTTTAAATTAAAAATGAAGTTTTAAATCAATCTAAAGTATATATGAGTAAACTTGGTCTGACAGTTACCAATGCTTAATCAGTGAGGCACCTATCTCAGCGATCTGTCTATTTCGTTCATCCATAGTTGCCTGACTCCCCGTCGTGTAGATAACTACGATACGGGAGGGCTTACCATCTGGCCCCAGTGCTGCAATGATACCGCGAGACCCACGCTCACCGGCTCCAGATTTATCAGCAATAAACCAGCCAGCCGGAAGGGCCGAGCGCAGAAGTGGTCCTGCAACTTTATCCGCCTCCATCCAGTCTATTAATTGTTGCCGGGAAGCTAGAGTAAGTAGTTCGCCAGTTAATAGTTTGCGCAACGTTGTTGCCATTGCTACAGGCATCGTGGTGTCACGCTCGTCGTTTGGTATGGCTTCATTCAGCTCCGGTTCCCAACGATCAAGGCGAGTTACATGATCCCCCATGTTGTGCAAAAAAGCGGTTAGCTCCTTCGGTCCTCCGATCGTTGTCAGAAGTAAGTTGGCCGCAGTGTTATCACTCATGGTTATGGCAGCACTGCATAATTCTCTTACTGTCATGCCATCCGTAAGATGCTTTTCTGTGACTGGTGAGTACTCAACCAAGTCATTCTGAGAATAGTGTATGCGGCGACCGAGTTGCTCTTGCCCGGCGTCAATACGGGATAATACCGCGCCACATAGCAGAACTTTAAAAGTGCTCATCATTGGAAAACGTTCTTCGGGGCGAAAACTCTCAAGGATCTTACCGCTGTTGAGATCCAGTTCGATGTAACCCACTCGTGCACCCAACTGATCTTCAGCATCTTTTACTTTCACCAGCGTTTCTGGGTGAGCAAAAACAGGAAGGCAAAATGCCGCAAAAAAGGGAATAAGGGCGACACGGAAATGTTGAATACTCATACTCTTCCTTTTTCAATATTATTGAAGCATTTATCAGGGTTATTGTCTCATGAGCGGATACATATTTGAATGTATTTAGAAAAATAAACAAATAGGGGTTCCGCGCACATTTCCCCGAAAAGTGCCACCTGACGTCTAAGAAACCATTATTATCATGACATTAACCTATAAAAATAGGCGTATCACGAGGCCCTTTCGTCTCGCGCGTTTCGGTGATGACGGTGAAAACCTCTGACACATGCAGCTCCCGGAGACGGTCACAGCTTGTCTGTAAGCGGATGCCGGGAGCAGACAAGCCCGTCAGGGCGCGTCAGCGGGTGTTGGCGGGTGTCGGGGCTGGCTTAACTATGCGGCATCAGAGCAGATTGTACTGAGAGTGCACCATATGCGGTGTGAAATACCGCACAGATGCGTAAGGAGAAAATACCGCATCAGGCGCCATTCGCCATTCAGGCTGCGCAACTGTTGGGAAGGGCGATCGGTGCGGGCCTCTTCGCTATTACGCCAGCTGGCGAAAGGGGGATGTGCTGCAAGGCGATTAAGTTGGGTAACGCCAGGGTTTTCCCAGTCACGACGTTGTAAAACGACGGCCAGTGCCAAGCTG | For generating MERVL-tdTomato mESC reporter cell line |
| pCas9-Gata2-2XsgRNA | gagggcctatttcccatgattccttcatatttgcatatacgatacaaggctgttagagagataattggaattaatttgactgtaaacacaaagatattagtacaaaatacgtgacgtagaaagtaataatttcttgggtagtttgcagttttaaaattatgttttaaaatggactatcatatgcttaccgtaacttgaaagtatttcgatttcttggctttatatatcttgtggaaaggacgaaacacccctgggctgtgcaacaagtggttttagagctagaaatagcaagttaaaataaggctagtccgttatcaacttgaaaaagtggcaccgagtcggtgcttttttgttttagagctagaaatagcaagttaaaataaggctagtccgtttttagcgcgtgcgccaattctgcagacaaatggctctagagagggcctatttcccatgattccttcatatttgcatatacgatacaaggctgttagagagataattggaattaatttgactgtaaacacaaagatattagtacaaaatacgtgacgtagaaagtaataatttcttgggtagtttgcagttttaaaattatgttttaaaatggactatcatatgcttaccgtaacttgaaagtatttcgatttcttggctttatatatcttgtggaaaggacgaaacaccgctgccatagtcatgagctggttttagagctagaaatagcaagttaaaataaggctagtccgttatcaacttgaaaaagtggcaccgagtcggtgcttttttgttttagagctagaaatagcaagttaaaataaggctagtccgtttttagcgcgtgcgccaattctgcagacaaatggggtacccgttacataacttacggtaaatggcccgcctggctgaccgcccaacgacccccgcccattgacgtcaatagtaacgccaatagggactttccattgacgtcaatgggtggagtatttacggtaaactgcccacttggcagtacatcaagtgtatcatatgccaagtacgccccctattgacgtcaatgacggtaaatggcccgcctggcattgtgcccagtacatgaccttatgggactttcctacttggcagtacatctacgtattagtcatcgctattaccatggtcgaggtgagccccacgttctgcttcactctccccatctcccccccctccccacccccaattttgtatttatttattttttaattattttgtgcagcgatgggggcggggggggggggggggcgcgcgccaggcggggcggggcggggcgaggggcggggcggggcgaggcggagaggtgcggcggcagccaatcagagcggcgcgctccgaaagtttccttttatggcgaggcggcggcggcggcggccctataaaaagcgaagcgcgcggcgggcgggagtcgctgcgcgctgccttcgccccgtgccccgctccgccgccgcctcgcgccgcccgccccggctctgactgaccgcgttactcccacaggtgagcgggcgggacggcccttctcctccgggctgtaattagctgagcaagaggtaagggtttaagggatggttggttggtggggtattaatgtttaattacctggagcacctgcctgaaatcactttttttcaggttggaccggtgccaccatggactataaggaccacgacggagactacaaggatcatgatattgattacaaagacgatgacgataagatggccccaaagaagaagcggaaggtcggtatccacggagtcccagcagccgacaagaagtacagcatcggcctggacatcggcaccaactctgtgggctgggccgtgatcaccgacgagtacaaggtgcccagcaagaaattcaaggtgctgggcaacaccgaccggcacagcatcaagaagaacctgatcggagccctgctgttcgacagcggcgaaacagccgaggccacccggctgaagagaaccgccagaagaagatacaccagacggaagaaccggatctgctatctgcaagagatcttcagcaacgagatggccaaggtggacgacagcttcttccacagactggaagagtccttcctggtggaagaggataagaagcacgagcggcaccccatcttcggcaacatcgtggacgaggtggcctaccacgagaagtaccccaccatctaccacctgagaaagaaactggtggacagcaccgacaaggccgacctgcggctgatctatctggccctggcccacatgatcaagttccggggccacttcctgatcgagggcgacctgaaccccgacaacagcgacgtggacaagctgttcatccagctggtgcagacctacaaccagctgttcgaggaaaaccccatcaacgccagcggcgtggacgccaaggccatcctgtctgccagactgagcaagagcagacggctggaaaatctgatcgcccagctgcccggcgagaagaagaatggcctgttcggaaacctgattgccctgagcctgggcctgacccccaacttcaagagcaacttcgacctggccgaggatgccaaactgcagctgagcaaggacacctacgacgacgacctggacaacctgctggcccagatcggcgaccagtacgccgacctgtttctggccgccaagaacctgtccgacgccatcctgctgagcgacatcctgagagtgaacaccgagatcaccaaggcccccctgagcgcctctatgatcaagagatacgacgagcaccaccaggacctgaccctgctgaaagctctcgtgcggcagcagctgcctgagaagtacaaagagattttcttcgaccagagcaagaacggctacgccggctacattgacggcggagccagccaggaagagttctacaagttcatcaagcccatcctggaaaagatggacggcaccgaggaactgctcgtgaagctgaacagagaggacctgctgcggaagcagcggaccttcgacaacggcagcatcccccaccagatccacctgggagagctgcacgccattctgcggcggcaggaagatttttacccattcctgaaggacaaccgggaaaagatcgagaagatcctgaccttccgcatcccctactacgtgggccctctggccaggggaaacagcagattcgcctggatgaccagaaagagcgaggaaaccatcaccccctggaacttcgaggaagtggtggacaagggcgcttccgcccagagcttcatcgagcggatgaccaacttcgataagaacctgcccaacgagaaggtgctgcccaagcacagcctgctgtacgagtacttcaccgtgtataacgagctgaccaaagtgaaatacgtgaccgagggaatgagaaagcccgccttcctgagcggcgagcagaaaaaggccatcgtggacctgctgttcaagaccaaccggaaagtgaccgtgaagcagctgaaagaggactacttcaagaaaatcgagtgcttcgactccgtggaaatctccggcgtggaagatcggttcaacgcctccctgggcacataccacgatctgctgaaaattatcaaggacaaggacttcctggacaatgaggaaaacgaggacattctggaagatatcgtgctgaccctgacactgtttgaggacagagagatgatcgaggaacggctgaaaacctatgcccacctgttcgacgacaaagtgatgaagcagctgaagcggcggagatacaccggctggggcaggctgagccggaagctgatcaacggcatccgggacaagcagtccggcaagacaatcctggatttcctgaagtccgacggcttcgccaacagaaacttcatgcagctgatccacgacgacagcctgacctttaaagaggacatccagaaagcccaggtgtccggccagggcgatagcctgcacgagcacattgccaatctggccggcagccccgccattaagaagggcatcctgcagacagtgaaggtggtggacgagctcgtgaaagtgatgggccggcacaagcccgagaacatcgtgatcgaaatggccagagagaaccagaccacccagaagggacagaagaacagccgcgagagaatgaagcggatcgaagagggcatcaaagagctgggcagccagatcctgaaagaacaccccgtggaaaacacccagctgcagaacgagaagctgtacctgtactacctgcagaatgggcgggatatgtacgtggaccaggaactggacatcaaccggctgtccgactacgatgtggaccatatcgtgcctcagagctttctgaaggacgactccatcgacaacaaggtgctgaccagaagcgacaagaaccggggcaagagcgacaacgtgccctccgaagaggtcgtgaagaagatgaagaactactggcggcagctgctgaacgccaagctgattacccagagaaagttcgacaatctgaccaaggccgagagaggcggcctgagcgaactggataaggccggcttcatcaagagacagctggtggaaacccggcagatcacaaagcacgtggcacagatcctggactcccggatgaacactaagtacgacgagaatgacaagctgatccgggaagtgaaagtgatcaccctgaagtccaagctggtgtccgatttccggaaggatttccagttttacaaagtgcgcgagatcaacaactaccaccacgcccacgacgcctacctgaacgccgtcgtgggaaccgccctgatcaaaaagtaccctaagctggaaagcgagttcgtgtacggcgactacaaggtgtacgacgtgcggaagatgatcgccaagagcgagcaggaaatcggcaaggctaccgccaagtacttcttctacagcaacatcatgaactttttcaagaccgagattaccctggccaacggcgagatccggaagcggcctctgatcgagacaaacggcgaaaccggggagatcgtgtgggataagggccgggattttgccaccgtgcggaaagtgctgagcatgccccaagtgaatatcgtgaaaaagaccgaggtgcagacaggcggcttcagcaaagagtctatcctgcccaagaggaacagcgataagctgatcgccagaaagaaggactgggaccctaagaagtacggcggcttcgacagccccaccgtggcctattctgtgctggtggtggccaaagtggaaaagggcaagtccaagaaactgaagagtgtgaaagagctgctggggatcaccatcatggaaagaagcagcttcgagaagaatcccatcgactttctggaagccaagggctacaaagaagtgaaaaaggacctgatcatcaagctgcctaagtactccctgttcgagctggaaaacggccggaagagaatgctggcctctgccggcgaactgcagaagggaaacgaactggccctgccctccaaatatgtgaacttcctgtacctggccagccactatgagaagctgaagggctcccccgaggataatgagcagaaacagctgtttgtggaacagcacaagcactacctggacgagatcatcgagcagatcagcgagttctccaagagagtgatcctggccgacgctaatctggacaaagtgctgtccgcctacaacaagcaccgggataagcccatcagagagcaggccgagaatatcatccacctgtttaccctgaccaatctgggagcccctgccgccttcaagtactttgacaccaccatcgaccggaagaggtacaccagcaccaaagaggtgctggacgccaccctgatccaccagagcatcaccggcctgtacgagacacggatcgacctgtctcagctgggaggcgacaaaaggccggcggccacgaaaaaggccggccaggcaaaaaagaaaaagtaagaattcctagagctcgctgatcagcctcgactgtgccttctagttgccagccatctgttgtttgcccctcccccgtgccttccttgaccctggaaggtgccactcccactgtcctttcctaataaaatgaggaaattgcatcgcattgtctgagtaggtgtcattctattctggggggtggggtggggcaggacagcaagggggaggattgggaagagaatagcaggcatgctggggagcggccgcaggaacccctagtgatggagttggccactccctctctgcgcgctcgctcgctcactgaggccgggcgaccaaaggtcgcccgacgcccgggctttgcccgggcggcctcagtgagcgagcgagcgcgcagctgcctgcaggggcgcctgatgcggtattttctccttacgcatctgtgcggtatttcacaccgcatacgtcaaagcaaccatagtacgcgccctgtagcggcgcattaagcgcggcgggtgtggtggttacgcgcagcgtgaccgctacacttgccagcgccttagcgcccgctcctttcgctttcttcccttcctttctcgccacgttcgccggctttccccgtcaagctctaaatcgggggctccctttagggttccgatttagtgctttacggcacctcgaccccaaaaaacttgatttgggtgatggttcacgtagtgggccatcgccctgatagacggtttttcgccctttgacgttggagtccacgttctttaatagtggactcttgttccaaactggaacaacactcaactctatctcgggctattcttttgatttataagggattttgccgatttcggtctattggttaaaaaatgagctgatttaacaaaaatttaacgcgaattttaacaaaatattaacgtttacaattttatggtgcactctcagtacaatctgctctgatgccgcatagttaagccagccccgacacccgccaacacccgctgacgcgccctgacgggcttgtctgctcccggcatccgcttacagacaagctgtgaccgtctccgggagctgcatgtgtcagaggttttcaccgtcatcaccgaaacgcgcgagacgaaagggcctcgtgatacgcctatttttataggttaatgtcatgataataatggtttcttagacgtcaggtggcacttttcggggaaatgtgcgcggaacccctatttgtttatttttctaaatacattcaaatatgtatccgctcatgagacaataaccctgataaatgcttcaataatattgaaaaaggaagagtatgagtattcaacatttccgtgtcgcccttattcccttttttgcggcattttgccttcctgtttttgctcacccagaaacgctggtgaaagtaaaagatgctgaagatcagttgggtgcacgagtgggttacatcgaactggatctcaacagcggtaagatccttgagagttttcgccccgaagaacgttttccaatgatgagcacttttaaagttctgctatgtggcgcggtattatcccgtattgacgccgggcaagagcaactcggtcgccgcatacactattctcagaatgacttggttgagtactcaccagtcacagaaaagcatcttacggatggcatgacagtaagagaattatgcagtgctgccataaccatgagtgataacactgcggccaacttacttctgacaacgatcggaggaccgaaggagctaaccgcttttttgcacaacatgggggatcatgtaactcgccttgatcgttgggaaccggagctgaatgaagccataccaaacgacgagcgtgacaccacgatgcctgtagcaatggcaacaacgttgcgcaaactattaactggcgaactacttactctagcttcccggcaacaattaatagactggatggaggcggataaagttgcaggaccacttctgcgctcggcccttccggctggctggtttattgctgataaatctggagccggtgagcgtggaagccgcggtatcattgcagcactggggccagatggtaagccctcccgtatcgtagttatctacacgacggggagtcaggcaactatggatgaacgaaatagacagatcgctgagataggtgcctcactgattaagcattggtaactgtcagaccaagtttactcatatatactttagattgatttaaaacttcatttttaatttaaaaggatctaggtgaagatcctttttgataatctcatgaccaaaatcccttaacgtgagttttcgttccactgagcgtcagaccccgtagaaaagatcaaaggatcttcttgagatcctttttttctgcgcgtaatctgctgcttgcaaacaaaaaaaccaccgctaccagcggtggtttgtttgccggatcaagagctaccaactctttttccgaaggtaactggcttcagcagagcgcagataccaaatactgttcttctagtgtagccgtagttaggccaccacttcaagaactctgtagcaccgcctacatacctcgctctgctaatcctgttaccagtggctgctgccagtggcgataagtcgtgtcttaccgggttggactcaagacgatagttaccggataaggcgcagcggtcgggctgaacggggggttcgtgcacacagcccagcttggagcgaacgacctacaccgaactgagatacctacagcgtgagctatgagaaagcgccacgcttcccgaagggagaaaggcggacaggtatccggtaagcggcagggtcggaacaggagagcgcacgagggagcttccagggggaaacgcctggtatctttatagtcctgtcgggtttcgccacctctgacttgagcgtcgatttttgtgatgctcgtcaggggggcggagcctatggaaaaacgccagcaacgcggcctttttacggttcctggccttttgctggccttttgctcacatgcgccattctcgagcatgcgccattctcgagcatgctcgagaatggcgcatgt | For generating Gata2 KO mESC cell line |
| pTS-EBFP-G4^Gata2^-DsRed | acatttgcttctgacacaactgtgttcactagcaacctcaaacagacGCCACCATGGTGAGCAAGGGCGAGGAGCTGTTCACCGGGGTGGTGCCCATCCTGGTCGAGCTGGACGGCGACGTAAACGGCCACAAGTTCAGCGTGTCCGGCGAGGGCGAGGGCGATGCCACCTACGGCAAGCTGACCCTGAAGTTCATCTGCACCACCGGCAAGCTGCCCGTGCCCTGGCCCACCCTCGTGACCACCCTGACCCACGGCGTGCAGTGCTTCAGCCGCTACCCCGACCACATGAAGCAGCACGACTTCTTCAAGTCCGCCATGCCCGAAGGCTACGTCCAGGAGCGCACCATCTTCTTCAAGGACGACGGCAACTACAAGACCCGCGCCGAGGTGAAGTTCGAGGGCGACACCCTGGTGAACCGCATCGAGCTGAAGGGCATCGACTTCAAGGAGGACGGCAACATCCTGGGGCACAAGCTGGAGTACAACTTCAACAGCCACAACGTCTATATCATGGCCGACAAGCAGAAGAACGGCATCAAGGTGAACTTCAAGATCCGCCACAACATCGAGGACGGCAGCGTGCAGCTCGCCGACCACTACCAGCAGAACACCCCCATCGGCGACGGCCCCGTGCTGCTGCCCGACAACCACTACCTGAGCACCCAGTCCGCCCTGAGCAAAGACCCCAACGAGAAGCGCGATCACATGGTCCTGCTGGAGTTCGTGACCGCCGCCGGGATCACTCTCGGCATGGACGAGCTGTACAAGGGATCTGGCGCCACCAACTTCTCTCTGCTGAAGCAGGCCGGCGACGTGGAGGAGAACCCAGGCCCAACTAGTGAGGTGGCGCCTGAGCAGCCTCGCTGGATGGCGCACCCCGCCGTATTGAATGCGCAGCACCCCGACTCGCACCATCCGGGCCTGGCGCATAACTACATGGAGCCAGCACAGCTGCTGCCTCCCGACGAGGTGGATGTCTTCTTCAACCATCTCGACTCGCAGGGCAACCCTTACTACGCCAACCCGGCCCACGCGCGCGCGCGCGTTTCCTACAGCCCGGCGCATGCCCGTCTCACCGGAGGCCAGATGTGCCGACCACACTTGTTGCACAGCCCAGGCTTGCCGTGGCTGGACGGGGGCAAAGCAGCTCTCTCTGCCGCCGCTGCCCATCACCACAGTCCCTGGACCGTCAGCCCGTTCTCCAAGACCCCGCTGCACCCCTCAGCTGCTGGAGCACCCGGAGGGCCTCTGTCTGTTTACCCAGGGGCTGCGGGTGGGAGCGGGGGAGGCAGTGGGAGCTCCGATAGCACTGAGAGCGGCTCCACCGAGTCCGTCATCAAGGAGTTCATGCGCTTCAAGGTGCACATGGAGGGCTCCGTGAACGGCCACGAGTTCGAGATCGAGGGCGAGGGCGAGGGCCGCCCCTACGAGGGCACCCAGACCGCCAAGCTGAAGGTGACCAAGGGTGGCCCCCTGCCCTTCGCCTGGGACATCCTGTCCCCTCAGTTCATGTACGGCTCCAAGGCCTACGTGAAGCACCCCGCCGACATCCCCGACTACTTGAAGCTGTCCTTCCCCGAGGGCTTCAAGTGGGAGCGCGTGATGAACTTCGAGGACGGCGGCGTGGTGACCGTGACCCAGGACTCCTCCCTGCAGGACGGCGAGTTCATCTACAAGGTGAAGCTGCGCGGCACCAACTTCCCCTCCGACGGCCCCGTAATGCAGAAGAAGACTATGGGCTGGGAGGCCTCCTCCGAGCGGATGTACCCCGAGGACGGCGCCCTGAAGGGCGAGATCAAGCAGAGGCTGAAGCTGAAGGACGGCGGCCACTACGACGCTGAGGTCAAGACCACCTACAAGGCCAAGAAGCCCGTGCAGCTGCCCGGCGCCTACAACGTCAACATCAAGTTGGACATCACCTCCCACAACGAGGACTACACCATCGTGGAACAGTACGAACGCGCCGAGGGCCGCCACTCCGGCTCCCAGGGAGGAAGCGGAGGAAGTTAAgtttaaagctcgctttcttgctgtccaatttctattaaaggttcctttgttccctaagtccaactactaaactgggggatattatgaagggccttgagcatctggattctgcctaataaaaaacatttattttcattgcaaAAAAAAAAAAAAAAAAAAAAAAAAAAAAAAaGCATATGACTAAAAAAAAAAAAAAAAAAAAAAAAAAAAAAAAAAAAAAAAAAAAAAAAAAAAAAAAAAAAAAAAAAAAAA | Encoding EBFP-G4^Gata2^-DsRed mRNA |
| pTS-EBFP-G4^Gata2-MUT^-DsRed | acatttgcttctgacacaactgtgttcactagcaacctcaaacagacGCCACCATGGTGAGCAAGGGCGAGGAGCTGTTCACCGGGGTGGTGCCCATCCTGGTCGAGCTGGACGGCGACGTAAACGGCCACAAGTTCAGCGTGTCCGGCGAGGGCGAGGGCGATGCCACCTACGGCAAGCTGACCCTGAAGTTCATCTGCACCACCGGCAAGCTGCCCGTGCCCTGGCCCACCCTCGTGACCACCCTGACCCACGGCGTGCAGTGCTTCAGCCGCTACCCCGACCACATGAAGCAGCACGACTTCTTCAAGTCCGCCATGCCCGAAGGCTACGTCCAGGAGCGCACCATCTTCTTCAAGGACGACGGCAACTACAAGACCCGCGCCGAGGTGAAGTTCGAGGGCGACACCCTGGTGAACCGCATCGAGCTGAAGGGCATCGACTTCAAGGAGGACGGCAACATCCTGGGGCACAAGCTGGAGTACAACTTCAACAGCCACAACGTCTATATCATGGCCGACAAGCAGAAGAACGGCATCAAGGTGAACTTCAAGATCCGCCACAACATCGAGGACGGCAGCGTGCAGCTCGCCGACCACTACCAGCAGAACACCCCCATCGGCGACGGCCCCGTGCTGCTGCCCGACAACCACTACCTGAGCACCCAGTCCGCCCTGAGCAAAGACCCCAACGAGAAGCGCGATCACATGGTCCTGCTGGAGTTCGTGACCGCCGCCGGGATCACTCTCGGCATGGACGAGCTGTACAAGGGATCTGGCGCCACCAACTTCTCTCTGCTGAAGCAGGCCGGCGACGTGGAGGAGAACCCAGGCCCAACTAGTGAGGTGGCGCCTGAGCAGCCTCGCTGGATGGCGCACCCCGCCGTATTGAATGCGCAGCACCCCGACTCGCACCATCCGGGCCTGGCGCATAACTACATGGAGCCAGCACAGCTGCTGCCTCCCGACGAGGTGGATGTCTTCTTCAACCATCTCGACTCGCAGGGCAACCCTTACTACGCCAACCCGGCCCACGCGCGCGCGCGCGTTTCCTACAGCCCGGCGCATGCCCGTCTCACCGGAGGCCAGATGTGCCGACCACACTTGTTGCACAGCCCAGGCTTGCCGTGGCTGGACGGGGGCAAAGCAGCTCTCTCTGCCGCCGCTGCCCATCACCACAGTCCCTGGACCGTCAGCCCGTTCTCCAAGACCCCGCTGCACCCCTCAGCTGCTGGAGCACCCGGAGGGCCTCTGTCTGTTTACCCAGGAGCTGCAGGTGGAAGCGGAGGAGGCTCCGGATCCTCCGATAGCACTGAGAGCGGCTCCACCGAGTCCGTCATCAAGGAGTTCATGCGCTTCAAGGTGCACATGGAGGGCTCCGTGAACGGCCACGAGTTCGAGATCGAGGGCGAGGGCGAGGGCCGCCCCTACGAGGGCACCCAGACCGCCAAGCTGAAGGTGACCAAGGGTGGCCCCCTGCCCTTCGCCTGGGACATCCTGTCCCCTCAGTTCATGTACGGCTCCAAGGCCTACGTGAAGCACCCCGCCGACATCCCCGACTACTTGAAGCTGTCCTTCCCCGAGGGCTTCAAGTGGGAGCGCGTGATGAACTTCGAGGACGGCGGCGTGGTGACCGTGACCCAGGACTCCTCCCTGCAGGACGGCGAGTTCATCTACAAGGTGAAGCTGCGCGGCACCAACTTCCCCTCCGACGGCCCCGTAATGCAGAAGAAGACTATGGGCTGGGAGGCCTCCTCCGAGCGGATGTACCCCGAGGACGGCGCCCTGAAGGGCGAGATCAAGCAGAGGCTGAAGCTGAAGGACGGCGGCCACTACGACGCTGAGGTCAAGACCACCTACAAGGCCAAGAAGCCCGTGCAGCTGCCCGGCGCCTACAACGTCAACATCAAGTTGGACATCACCTCCCACAACGAGGACTACACCATCGTGGAACAGTACGAACGCGCCGAGGGCCGCCACTCCGGCTCCCAGGGAGGAAGCGGAGGAAGTTAA | Encoding EBFP-G4^Gata2-MUT^-DsRed mRNA |

**Table S3. Oligos used in this work.**

| **Oligonucleotide** | **Sequence** | **Note** |
| --- | --- | --- |
| Gapdh-F | GACATCAAGAAGGTGGTGAAGC | RT‒qPCR primers |
| Gapdh-R | ACCCTGTTGCTGTAGCCGTA |  |
| Zscan4d-F | TGAGGTGGAGGAGTAGGTAAAC |  |
| Zscan4d-R | ACCAGGAGTTGAACATCTTCC |  |
| Zscan4f-F | AATGAGCAGATGCCAGTAGACA |  |
| Zscan4f-R | GATAAATCCTTTGGTTGTTCCTCC |  |
| Btg2-F | GGGAAGAGAACCGACATGCT |  |
| Btg2-R | CAGTGGTGTTTGTAATGATCGGT |  |
| Mdm2-F | ATCAGACAGGAGAAAGCGATACA |  |
| Mdm2-R | ATCCAAGCCTTCTTCTGCCT |  |
| Sp110-F | ATGAAGGTGAACATCGCCTATG |  |
| Sp110-R | GGACAGAGGGACCAGATTTTG |  |
| Cdx2-F | AGGCTGAGCCATGAGGAGTA |  |
| Cdx2-R | CGAGGTCCATAATTCCACTCA |  |
| Esx1-F | ATGGAATCTCACAAAAAGTGCCC |  |
| Esx1-R | ACTGCTCCAACAAAGGTCTTC |  |
| Tfap2c-F | AAGCGGTGGCTGACTATTTAA |  |
| Tfap2c-R | CAGGCTGAAATGAGACAAACAG |  |
| Elf5-F | ATTCGCTCGCAAGGTTACTCC |  |
| Elf5-R | GGATGCCACAGTTCTCTTCAG |  |
| Gata2-G4-A | AGGGGCUGCGGGUGGGAGCGGGGGA | RNA oligos for NMR |
| Gata2-G4-A-MUT1 | AGAAACUGCAAAUAAAAGCAAAGGA |  |
| Gata2-G4-A-MUT2 | AGGAGCUGCAGGUGGAAGCGGAGGA |  |
| Gata2-G4-B | UGGGAGCGGGGGAGGCAGUGGGA |  |
| Gata2-G4-B-MUT1 | UAAAAGCAAGAAAAGCAGUAAAA |  |
| Gata2-G4-B-MUT2 | UGGAAGCGGAGGAGGCUCCGGAA |  |
| hGata2-G4-A | AGGGGCUGGGGGUGGGAGCGGGGGA |  |
| hGata2-G4-A-MUT | AGAAGCUGAAAGUGAGAGCGAAGGA |  |
| hGata2-G4-B | UGGGGGUGGGAGCGGGGGAGGCAGCGGGA |  |
| hGata2-G4-B-MUT | UGAAAGUGAGAGCGAAAGAGACAGCGAGA |  |
| Gata2-G4-A truncated complementary sequence (TCS) | AGCCCC |  |
| Gata2-G4-B truncated complementary sequence (TCS) | CCCCCGCT | DNA oligos for NMR |
| hGata2-G4-A truncated complementary sequence (TCS) | CCCCCGCT |  |
| hGata2-G4-B truncated complementary sequence (TCS) | ACCCCC |  |

**Table S4. Published data used in this work.**

| **Deposited data** | **Source** | **Identifier** |
| --- | --- | --- |
| Zygote RNA-seq | NCBI Gene Expression Omnibus (GEO) | GSE66582 |
| 2-cell blastomere RNA-seq | NCBI Gene Expression Omnibus (GEO) | GSE66582 |
| Early 2-cell blastomere RNA-seq | NCBI Gene Expression Omnibus (GEO) | GSE66582 |
| Middle 2-cell blastomere RNA-seq | NCBI Gene Expression Omnibus (GEO) | GSE45719 |
| Late 2-cell blastomere RNA-seq | NCBI Gene Expression Omnibus (GEO) | GSE45719 |
| 4-cell blastomere RNA-seq | NCBI Gene Expression Omnibus (GEO) | GSE66582 |
| 8-cell blastomere RNA-seq | NCBI Gene Expression Omnibus (GEO) | GSE66582 |
| 16-cell blastomere RNA-seq | NCBI Gene Expression Omnibus (GEO) | GSE45719 |
| Early blastocyst RNA-seq | NCBI Gene Expression Omnibus (GEO) | GSE45719 |
| Middle blastocyst RNA-seq | NCBI Gene Expression Omnibus (GEO) | GSE45719 |
| Late blastocyst RNA-seq | NCBI Gene Expression Omnibus (GEO) | GSE45719 |
| E3.5 embryo RNA-seq | NCBI Gene Expression Omnibus (GEO) | GSE100597 |
| E4.5 embryo RNA-seq | NCBI Gene Expression Omnibus (GEO) | GSE100597 |
| E5.5 embryo RNA-seq | NCBI Gene Expression Omnibus (GEO) | GSE100597 |
| E6.5 embryo RNA-seq | NCBI Gene Expression Omnibus (GEO) | GSE100597 |
| mESC RNA-seq | NCBI Gene Expression Omnibus (GEO) | GSE66582 |
| CiTotiSC RNA-seq | NCBI Gene Expression Omnibus (GEO) | GSE185000 |
| D-EPSC RNA-seq | NCBI Gene Expression Omnibus (GEO) | GSE89303 |
| L-EPSC RNA-seq | NCBI Gene Expression Omnibus (GEO) | GSE145609 |
| miR-34a KO mESC RNA-seq | NCBI Gene Expression Omnibus (GEO) | GSE69484 |
| TBLC RNA-seq | NCBI Gene Expression Omnibus (GEO) | GSE168728 |
| TLSC RNA-seq | NCBI Gene Expression Omnibus (GEO) | GSE166216 |
| 2-cell like cell RNA-seq | NCBI Gene Expression Omnibus (GEO) | GSE33920 |
| 2-cell blastomere ATAC-seq | NCBI Gene Expression Omnibus (GEO) | GSE66581 |
| 4-cell blastomere ATAC-seq | NCBI Gene Expression Omnibus (GEO) | GSE66581 |
| 8-cell blastomere ATAC-seq | NCBI Gene Expression Omnibus (GEO) | GSE66581 |
| ICM ATAC-seq | NCBI Gene Expression Omnibus (GEO) | GSE66581 |
